# The Unkempt RNA binding protein reveals a local translation program in centriole overduplication

**DOI:** 10.1101/2024.07.29.605660

**Authors:** Abraham Martinez, Alexander J. Stemm-Wolf, Ryan M. Sheridan, Matthew J. Taliaferro, Chad G. Pearson

## Abstract

Excess centrosomes cause defects in mitosis, cell-signaling, and cell migration, and therefore their assembly is tightly regulated. Plk4 controls centriole duplication at the heart of centrosome assembly, and elevation of Plk4 promotes centrosome amplification (CA), a founding event of tumorigenesis. Here, we investigate the transcriptional consequences of elevated Plk4 and find Unkempt, a gene encoding an RNA binding protein with roles in translational regulation, to be one of only two upregulated mRNAs. Unk protein localizes to centrosomes and Cep131-positive centriolar satellites and is required for Plk4-induced centriole overduplication in an RNA-binding dependent manner. Translation is enriched at centrosomes and centriolar satellites with Unk and Cep131 promoting this localized translation. A transient centrosomal downregulation of translation occurs early in Plk4-induced CA. CNOT9, an Unk interactor and component of the translational inhibitory CCR4-NOT complex, localizes to centrosomes at this time. In summary, centriolar satellites and Unk promote local translation as part of a translational program that ensures centriole duplication.

## Introduction

Centrosomes, composed of a pair of centrioles and pericentriolar material, nucleate and organize the microtubules (MTs) of the bipolar mitotic spindle during cell division. During interphase, centrosomes along with the Golgi apparatus, nucleate MTs for general endo-lysosomal transport and for the transport of cargoes to and from the centrosome^1^. At G1 of the cell cycle, a single centrosome is present in each cell. A second centrosome is assembled during S phase and this is controlled primarily by the Polo-like kinase, Plk4, which phosphorylates and recruits proteins to the site of nascent centriole assembly^2,3^. Like DNA replication, centriole assembly is tightly regulated to once-and-only once each cell cycle^4^. The dysregulation of centriole assembly proteins can lead to centriole overduplication and centrosome amplification (CA), or the assembly of more than one centrosome. Consequences of CA include multipolar mitoses, cell death, aneuploidies, chromosome missegregation, and chromosomal instability^5–10^. These abnormalities result in human diseases, including microcephaly^11^ and cancer^12^. Therefore, the levels and spatial localization of centriole assembly proteins are tightly regulated to ensure that cells assemble only one centrosome during S phase.

Centriole duplication begins at the G1- to S-phase transition where Plk4 is recruited to the two centrioles of the centrosome. Plk4 associates with the centrioles through its interactions with the centriole wall complex proteins, Cep63-Cep152 and Cep192-Cep152^13,14^. Plk4 coalesces at the site of daughter, procentriole formation^15^. Plk4 phosphorylates STIL and promotes the recruitment of additional centriole proteins required for early stages of centriole assembly^2,15^. Following daughter centriole assembly, the two centrosomes comprising four centrioles mature through the cell cycle. Because the protein level and centrosome localization of Plk4 can dictate how many centrioles will form^2^, Plk4 is tightly regulated to prevent promiscuous centriole assembly^16–18^. An increase in the levels of Plk4 at centrioles promotes centriole overduplication and results in CA^5^. To prevent this, Plk4 auto-phosphorylation promotes its own degradation in a ubiquitin-mediated proteosome dependent manner^17^. Therefore, controlled centriole duplication and assembly requires that centriole proteins cooperate to regulate the stability of Plk4 and its specific localization at mother centrioles.

Plk4 also has a role in regulating its own stability at the centrioles through its effect on centriolar satellite proteins^19^. Centriolar satellites are membrane-less, electron dense granules that scaffold and transport cargoes to and from the centrosome in a MT dependent manner^20–22^. Cargoes include centrosome assembly proteins, ubiquitylating and deubiquitylating proteins, and components involved in ciliogenesis^23–26^. Plk4 phosphorylates the major scaffolding centriolar satellite proteins Cep131 and PCM1 to maintain their centrosomal localization, which in turn stabilizes Plk4 at centrioles^19^. Therefore, in addition to Plk4 regulating its own stability through phosphorylation, Plk4 relies on trafficking events for stabilization to promote centriole duplication.

Our current understanding of centriole duplication is limited to protein-protein interactions and the recruitment and transport to the centrosome of centriole assembly proteins that cooperate with Plk4. Recently, studies found mRNAs encoding centriole and centrosome proteins localize to centrosomes for local translation and centrosome regulation^27^. How RNAs are trafficked and the mechanisms by which translation is regulated at centrosomes is poorly understood. Local translation at the centrosome has been primarily studied in mitosis^28^, so whether local translation occurs at the centrosome to regulate centriole duplication is unknown. In support of this possibility, RNAs have been localized to the centrosome during interphase when centriole duplication and centrosome maturation occur^28^. Moreover, RNA binding proteins (RBPs) localize to centrosomes ^29–31^. Intriguingly, proteomics studies identifying interactions with centrosome and centriolar satellite proteins have found multiple RBPs, translation elongation and initiation factors, ribosomal proteins, and other RNA processing machinery^25,26,32^. This suggests that centriolar satellites may regulate centrosome-associated translation. Whether centriolar satellites are involved in the trafficking of RNAs, local translation, or other RNA regulatory processes concomitant with cell cycle dependent centrosome functions such as centriole duplication is unknown^24,33^. Because Plk4 phosphorylates centriolar satellite proteins to regulate their localization and cargo transport to the centrosome, centriolar satellite regulation may also be required for delivery of translational machinery, RNAs, or RNA processing complexes necessary for centriole duplication.

In this study, we identified the RBP, *Unkempt* (UNK) gene, to be one of few whose RNA is upregulated when Plk4 levels are elevated. Unk localizes to centrosomes and centriolar satellites, and its RNA binding activity is necessary for Plk4-induced centriole overduplication. Unk depletion decreases the amounts of centriole assembly proteins at centrosomes and results in the dispersal of the centriolar satellite proteins Cep131 and PCM1. Like Unk, Cep131, but not PCM1, promotes Plk4-induced centriole overduplication. We identify centriolar satellites as sites of active translation, and find that Unk, PCM1, and Cep131, promote the synthesis of proteins at the centrosome during late centriole assembly. We also find decreased translation at the centrosome early in centriole overduplication, suggesting that coordinating translation may be required for centriole overduplication. Finally, the Unk-interacting, mRNA silencing, and translation suppression CCR4-NOT complex subunits CNOT2 and CNOT9^34^ localize to centrioles, and the localization of CNOT9 increases at centrioles during early centriole assembly. These data suggest that Unk facilitates Plk4-induced centriole overduplication by positively regulating the localization of centriole assembly proteins, centriolar satellites, and translation at centrosomes.

## Results

### The Unkempt RNA binding protein facilitates Plk4-induced centriole overduplication

Elevated Plk4 is common in cancer cells^19,35^. Plk4 overexpression promotes the promiscuous assembly of multiple, nascent centrioles on the walls of mother centrioles (Fig. 1, A; ^36–38^). To determine whether a transcriptional response is associated with Plk4-induced centriole overduplication, we arrested RPE-1 cells in S phase and overexpressed Plk4 using a doxycycline inducible cell line^39^. RNA sequencing was performed on samples isolated 16 hours post Plk4 overexpression. Three transcripts were elevated by more than 1.5-fold: PLK4, UNK, and GALNT16. The *Unkempt* (UNK) RNA transcript was elevated similarly to overexpressed Plk4 (Fig. 1, B). The increase in UNK mRNA and its encoded protein in the Plk4 overexpressed cells was confirmed by fluorescence *in situ* hybridization (smFISH), western blot, and immunofluorescence (Fig. S1, A and B). Thus, Unk is upregulated in response to Plk4 overexpression.

**Figure 1.**
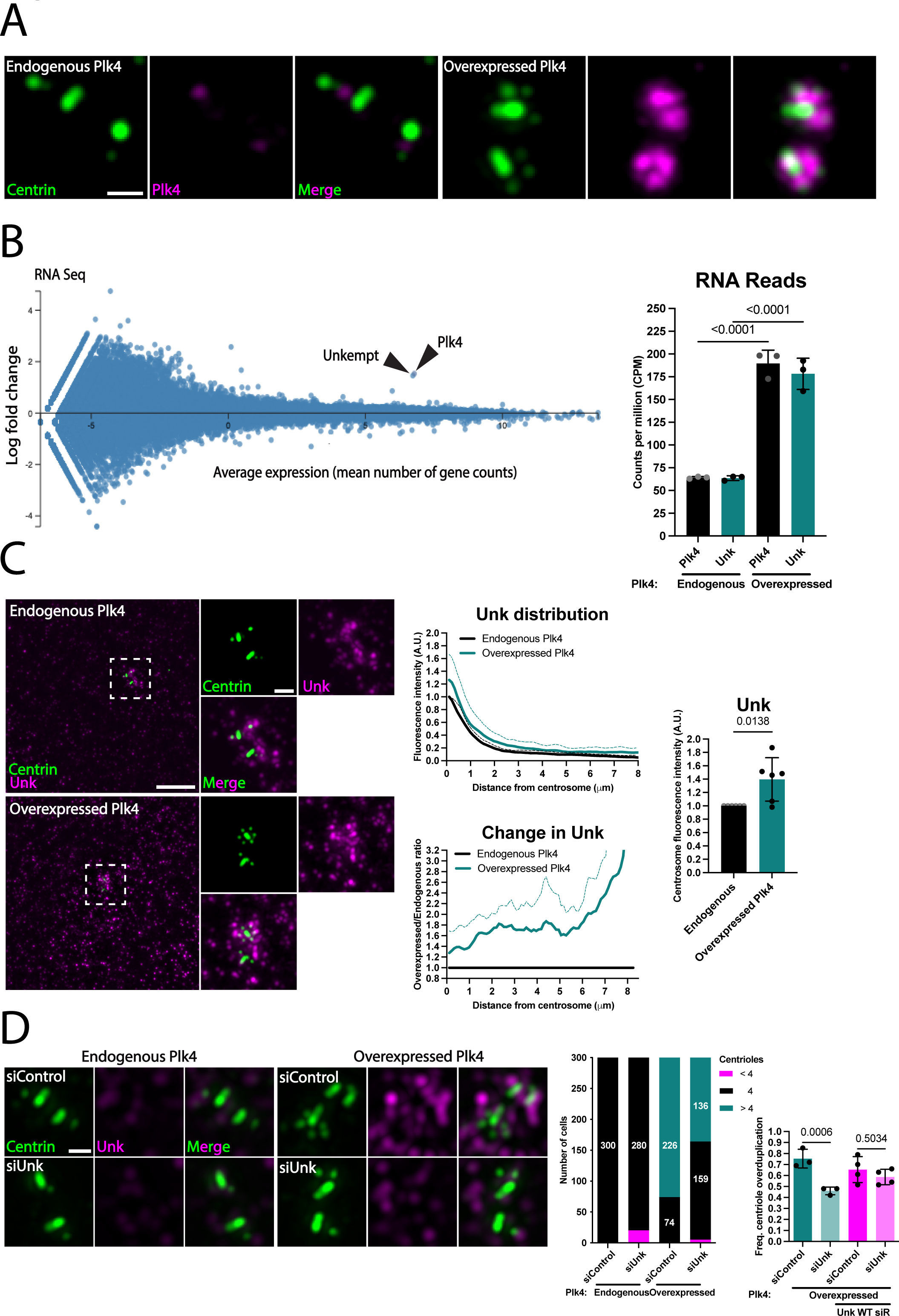
The Unkempt RNA binding protein facilitates Plk4-induced centriole overduplication. **(A)** Plk4 overexpression promotes centriole overduplication. Structured Illumination Microscopy (SIM) images of RPE-1 cells with endogenous and overexpressed Plk4. Plk4, magenta and centrioles, green. Scale bar 0.5 μm. **(B)** Unk is elevated upon Plk4 overexpression. Left panel – MA plot of mRNA levels based on RNA seq of RPE-1 cells with overexpressed Plk4, 16 hours after overexpression. PLK4 and UNK transcripts are elevated by 1.5- and 1.4-fold, respectively. Right panel – frequencies of PLK4 and UNK mRNAs in RPE-1 cells with endogenous and overexpressed Plk4. Graph values expressed as the means of biological replicates and SD. P values were determined using one-way ANOVA with Šídák *post hoc* test. **(C)** Unk protein is elevated at the centrosome by Plk4 overexpression. Left panels – SIM images of endogenous Unk in RPE-1 cells with endogenous and overexpressed Plk4. Unk, magenta and centrioles, green. Scale bar, 5 μm. Insets scale bar, 1 μm. Middle panels – 8 μm radial fluorescence intensity and corresponding ratio quantification of Unk using Centrin as the centrosome centroid. Right panel – centrosomal Unk fluorescence intensity based on binned central 2 μm. Graph values expressed as the means of biological replicates and SD. P value was determined using an unpaired two-tailed t test. **(D)** Unk is required for Plk4-induced centriole overduplication. Left panels – SIM images of RPE-1 cells showing centrioles in Unk depleted cells with endogenous and overexpressed Plk4. Unk, magenta and centrioles, green. Scale bar, 0.5 μm. Middle panel – frequency of cells with centriole overduplication. Right panel – number of cells with greater than, equal to, and less than four centrioles. Graph values expressed as the means of biological replicates and SD. P values were determined using one-way ANOVA with Šídák *post hoc* test.

To determine if Unkempt protein (Unk) functions at centrosomes, we visualized its subcellular localization. While the majority of Unk protein is cytoplasmic^40,41^, Unk also localized to the centrosome (Fig. 1, C). Further, co-localization of Unk with Centrin-labeled centrioles indicated that Unk resides near the distal ends of procentrioles (Fig. 1C). Similarly, mCherry tagged Unk localized to centrosomes (Fig. S1, C). To quantify the localization of Unk at the centrosome, radial fluorescence intensity analysis, hereafter radial analysis, was performed for Unk in an 8 μm radius around the centrosome. The centroid of the centrosome was defined by the Centrin centriole marker. Both cytoplasmic and centrosome levels of Unk increased upon Plk4 overexpression (Figs. 1, C and S1, B). To quantify the change in Unk distribution upon Plk4 overexpression, Unk fluorescence after Plk4 overexpression was normalized as a ratio (ratio analysis) to Unk fluorescence at endogenous Plk4 levels. Unk increased at the centrosome and in the cytoplasm. The centrosome population of Unk, as defined by the radial 2 μm around the centrosome, increased by approximately 40% upon Plk4 overexpression (Fig. 1, C). Thus, Unk localizes to centrioles and centrosomes and this localization increases upon Plk4-induced centriole overduplication.

To determine whether Unk is required for Plk4-induced centriole overduplication, S phase arrested cells were depleted of Unk by RNA interference (RNAi). Unk depletion did not affect canonical centriole duplication in cells without Plk4-induced centriole duplication (Fig. 1D). However, Unk depletion reduced the frequency of cells with Plk4-induced centriole overduplication from 75% (siControl) to 45% (siUnk) (Fig. 1D). Depletion of Unk with a second siRNA, showed similar results (Fig. S1, D). A knockout *TP53* WT RPE-1 clonal cell line targeting the endogenous *Unkempt* locus using CRISPR/cas9 could not be generated, suggesting that Unk may regulate canonical centriole overduplication and upon loss of centrioles, RPE-1 cells undergo p53-mediated cell death^42^. Centriole overduplication was rescued by an siRNA-resistant mCherry-Unk in Unk siRNA treated cells, suggesting that Unk is required for Plk4-induced centriole overduplication. The lack of an effect on canonical duplication may be explained by the partial depletion of Unk using RNA interference (Unk was depleted by 45% in Plk4 overexpressed cells and 24% in endogenous Plk4 cells). After Unk knockdown in Plk4 overexpressing cells, Unk levels remained higher than endogenous levels without Plk4 overexpression (26%; Fig. S1, B). In summary, Unk facilitates Plk4-induced centriole overduplication.

### Unkempt promotes localization of centriole assembly factors

To elucidate how Unk facilitates Plk4-induced centriole overduplication, we asked whether Unk regulates the localization of centriole assembly proteins, including Plk4. Increased Plk4 localization to centrioles is required for centriole overduplication^5^. To determine whether Unk modulates Plk4 protein localization to centrosomes, S phase arrested cells were depleted for Unk, prior to Plk4 overexpression. Centrosome localization of Plk4 protein was reduced by approximately 35% in Unk knockdown (Fig. 2, A). However, total cellular levels of Plk4 were unchanged (Fig. S2, A). This suggests that Unk facilitates centriole overduplication by promoting the localization of Plk4 protein to centrosomes.

**Figure 2.**
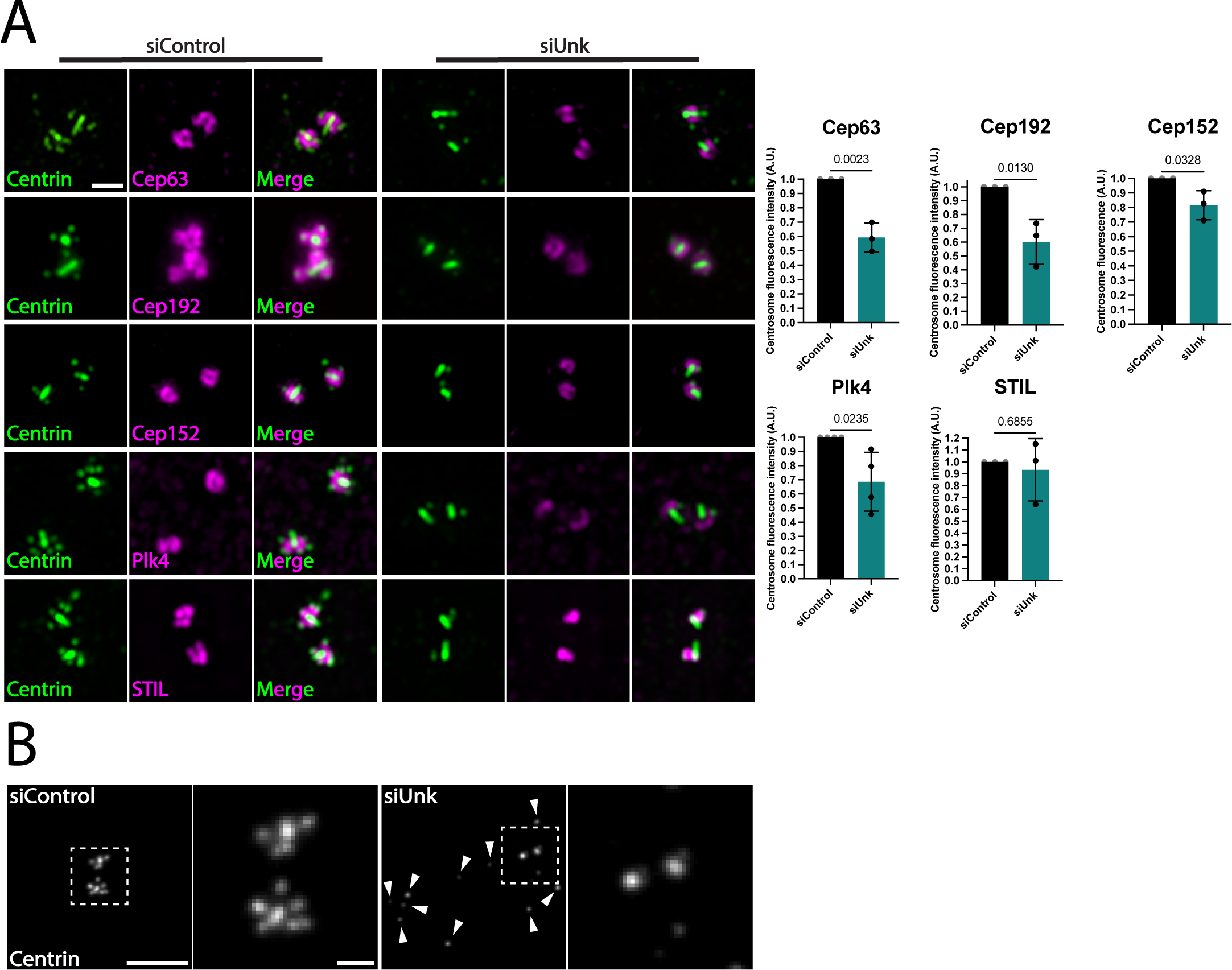
Unkempt promotes the localization of centriole assembly protein factors. **(A)** Centriole assembly protein localization is reduced in siUnk cells. Left panels – SIM images of RPE-1 cells showing centriolar fluorescence intensities at the centrosomes in Unk depleted cells with overexpressed Plk4. Cep63, Cep192, Cep152, Plk4, and STIL, magenta and centrioles, green. Scale bar, 1.0 μm. Right panels – average normalized centrosome fluorescence intensities of centriolar proteins. Graph values expressed as the means of biological replicates and SD. P values were determined using an unpaired two-tailed t test. **(B)** Unk depletion results in ectopic Centrin-positive foci. Confocal images of RPE-1 cells showing Centrin-positive foci in Unk depleted cells with overexpressed Plk4. Centrioles, greyscale. Arrows indicate sites of ectopic Centrin foci. Scale bars, Scale bar, 5 μm. Left insets scale bar, 0.5 μm. Right insets scale bar, 1 μm.

To determine how Unk promotes Plk4 centrosome localization, we asked whether Unk modulates centriole proteins that recruit and anchor Plk4. Cep63-Cep152, Cep192-Cep152 complexes, and STIL all recruit Plk4^13,14^. The mean centrosome fluorescence intensity of Cep63, Cep192, and Cep152 were all decreased upon knockdown of Unk (40%, 40%, and 20%, respectively; Fig. 2, A). A significant decrease in STIL was not observed (Fig. 2, A). This suggests that Unk promotes localization of early centriole assembly proteins required for Plk4 localization.

Centrin-positive puncta distributed throughout the cytoplasm were observed in Unk knockdown cells, regardless of Plk4 status (Fig. 2, B). These puncta were negative for other centriole (Plk4, Cep192, Cep152, or STIL), or centrosome (Pericentrin or CDK5RAP2) proteins, and did not nucleate microtubules (Fig. S2, B). This indicates that these puncta are not *bona fide* centrioles. The Centrin puncta were, however, positive for centriolar satellite proteins (PCM1 and Cep131) (Fig. S2, C). This suggests that these puncta are a subset of the centriolar satellites dispersed from the centrosome. We conclude that Unk promotes the localization of centriole assembly proteins required for centriole overduplication. Moreover, Unk depletion causes the dispersal of Centrin-positive puncta that co-stain with centriolar satellite proteins.

### Unkempt is a centriolar satellite protein and promotes localization of PCM1 and Cep131

Centriolar satellites are non-membranous granules surrounding centrosomes that transport proteins along MTs for centrosome assembly ^19,22,43^ and ciliogenesis^44–47^. Unk was found in a proximity labeling screen using multiple centriolar satellite protein baits^26^. We confirmed that Unk is a centriolar satellite protein by colocalization with the centriolar satellite marker, Cep131 (Fig. 3, A). Unk colocalizes with a subset of Cep131 cytoplasmic puncta and Cep131 puncta adjacent to procentrioles.

**Figure 3.**
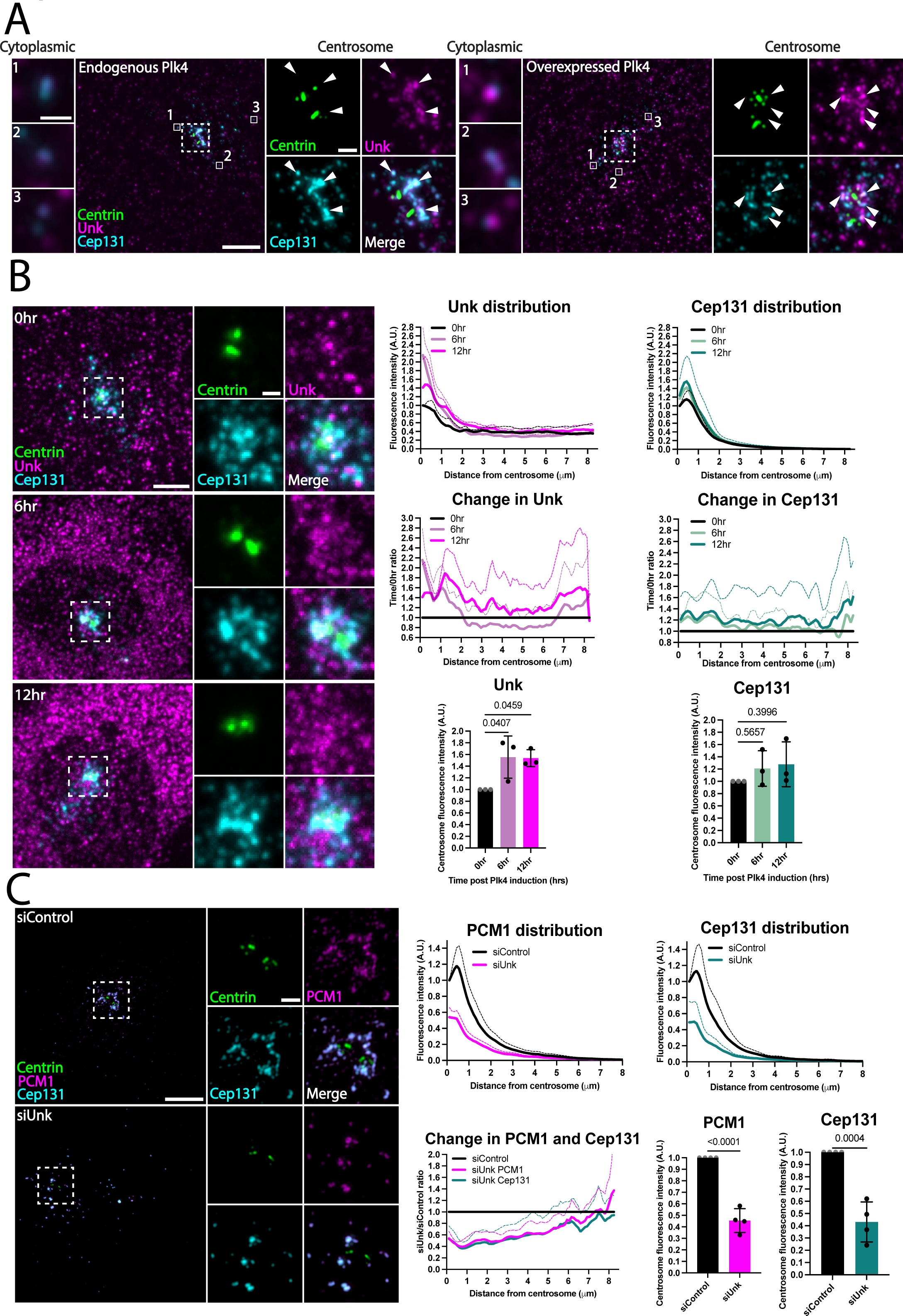
Unkempt is a centriolar satellite protein and promotes localization of PCM1 and Cep131. **(A)** Unk localizes to Cep131-positive centriolar satellites. SIM images of RPE-1 cells showing Unk protein localization with endogenous and overexpressed Plk4. Unk, magenta; Cep131, cyan; centrioles, green. Arrows indicate sites of Unk and Cep131 colocalization. Scale bar, 5 μm. Left insets scale bar, 0.5 μm. Right insets scale bar, 1 μm. **(B)** Plk4 overexpression promotes centriolar satellite and Unk localization to centrosomes. Left panels – confocal images of RPE-1 cells showing Unk and Cep131 localization 6 and 12 hours after Plk4 overexpression. Unk, magenta; Cep131, cyan; centrioles, green. Scale bar 5 μm. Insets scale bar, 1 μm. Right top and middle panels – 8 μm radial fluorescence intensity and corresponding ratio quantification of Unk and Cep131 using Centrin as the centrosome centroid. Right bottom - centrosomal Unk and Cep131 fluorescence intensity based on binned central 2 μm. Graph values expressed as the means of biological replicates and SD. P values were determined using one-way ANOVA with Dunnett *post hoc* test. **(C)** Centriolar satellites are reduced from the centrosome in siUnk cells with overexpressed Plk4. Left panels– SIM images of RPE-1 cells showing centriolar satellites and their localization to the centrosomes in Unk depleted cells with overexpressed Plk4. PCM1, magenta; Cep131, cyan; centrioles, green. Scale bar, 5 μm. Insets scale bar, 1.0 μm. Right, top panels – 8 μm radial fluorescence intensity of PCM1 and Cep131 using Centrin as the centrosome centroid. Right, bottom left panels - 8 μm ratio quantification of PCM1 and Cep131 using Centrin as the centroid. Right, bottom right panels - centrosomal PCM1 and Cep131 fluorescence intensity based on binned central 2 μm. Graph values expressed as the means of replicates and SD. P values were determined using an unpaired two-tailed t test.

We next asked how Cep131 and Unk localizations change during procentriole assembly. Centriole overduplication, indicated by Centrin-positive procentrioles, is observed 6 hours after Plk4 overexpression, and reached a maximum by 12 hours (Fig. S2, D). To capture the dynamics of Unk and Cep131, cells were analyzed 6 and 12 hours after Plk4 overexpression. Unk localization was specifically increased at the centrosome at 6 hours (45%). Although Unk levels were still increased 12 hours after Plk4 overexpression (45%), the localization was less specific to the centrosome (Fig 3, B). Cep131 exhibited a modest increase at the centrosome (20 and 25%) at both the 6 and 12 hour timepoints, respectively (Fig 3, B). Thus, Unk increases at centrosomes at early stages of Plk4-induced centriole overduplication, in a manner distinct from Cep131.

We next asked if Unk regulates centriolar satellites. Unk knockdown decreased centrosome localization of PCM1 and Cep131 centriolar satellite proteins causing dispersal throughout the cytoplasm (Fig. 3, C and S3, A). This redistribution of PCM1 and Cep131 upon Unk knockdown was independent of Plk4 overexpression (Fig. S3, A). The total cellular levels of PCM1 and Cep131 did not change upon Unk depletion (Fig. S3, B). Thus, Unk promotes the localization of centriolar satellites to the centrosome but does not affect the cellular levels of Cep131 (Fig. S3, A).

### Cep131, but not PCM1, promotes Plk4-induced centriole overduplication

Because Unk promotes centrosome localization of both PCM1- and Cep131-positive centriolar satellites (Fig. 3, C) and promotes Plk4-induced centriole overduplication (Fig. 1, D), we asked if these proteins themselves regulate Plk4-induced centriole overduplication. PCM1, Cep131, or both were knocked down in Plk4 overexpressing, S phase arrested cells. PCM1 knockdown did not affect the frequency of cells with overduplicated centrioles. In contrast, Cep131 knockdown and the double knockdown attenuated centriole overduplication to approximately 45% and 32%, respectively (Fig. 4, A). This is consistent with prior reports showing that loss of PCM1 does not affect centriole amplification in multiciliated cells^48^. Cep131 does suppress Plk4-induced centriole overduplication^19,22,48^. This suggests different roles for PCM1 and Cep131 in centriole assembly. Moreover, knockdown of PCM1 increased the centriole localization of Cep131 and Unk (not significant; Fig. 4, A, C), whereas knockdown of Cep131 decreased centrosome localization of PCM1^22,48^ and Unk (Fig. 4, A, C). In summary, knockdown of Cep131, but not PCM1, had a similar effect as Unk knockdown on Plk4-induced centriole overduplication. This suggests that Unk’s role in Plk4-induced centriole duplication may occur in coordination with Cep131, but not PCM1.

**Figure 4.**
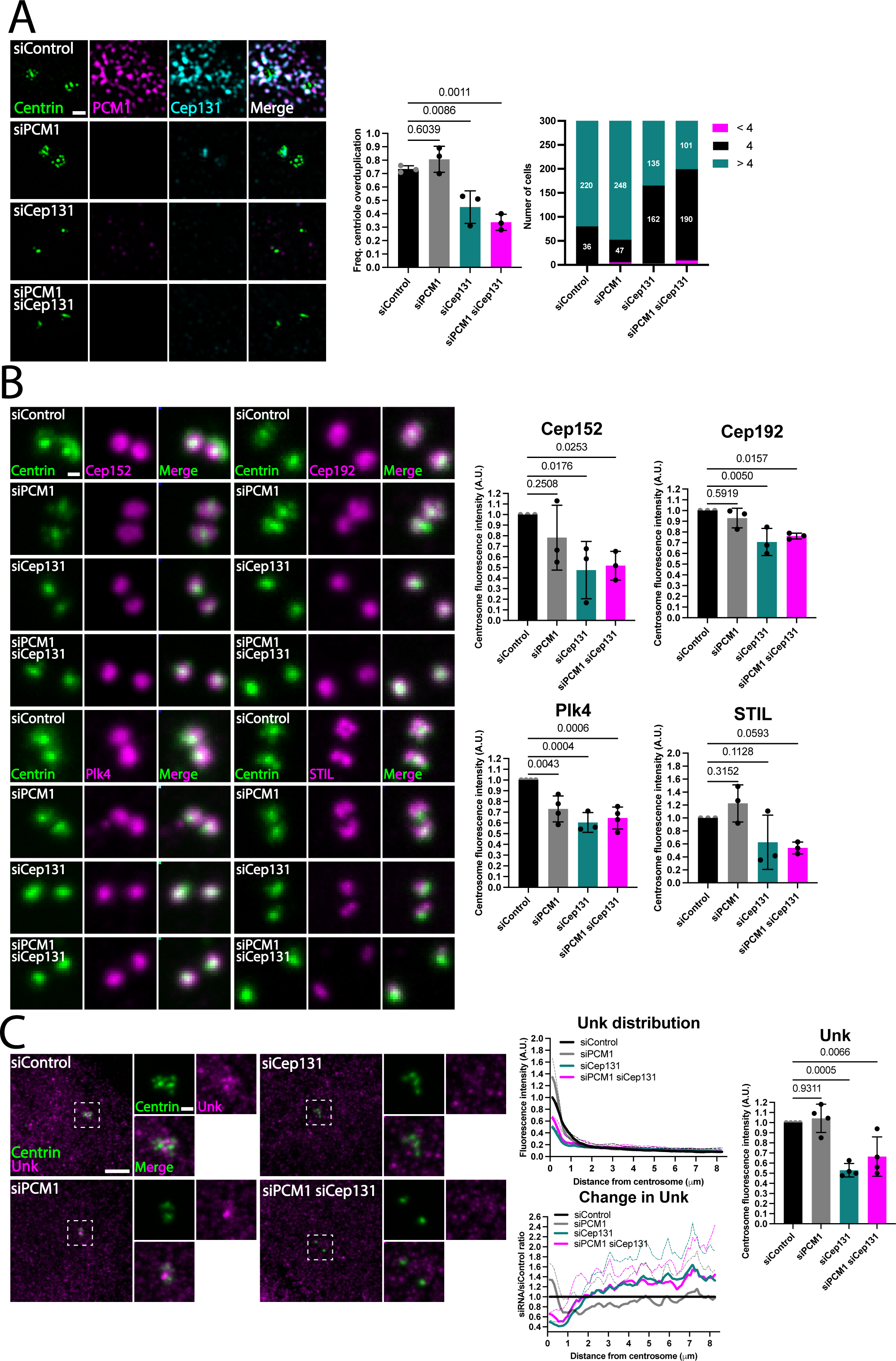
Cep131, but not PCM1, promotes Plk4-induced centriole overduplication. **(A)** Cep131 promotes Plk4-induced centriole overduplication. Left panels – SIM images of RPE-1 cells showing centrioles in PCM1 or Cep131, and double depleted cells with overexpressed Plk4. PCM1, magenta; Cep131, cyan; centrioles, green. Scale bar, 1.0 μm. Middle panel – frequency of cells with centriole overduplication. Right panel – number of cells with greater than, equal to, and less than 4 centrioles. Graph values expressed as the means of biological replicates and SD. P values were determined using one-way ANOVA with Dunnett *post hoc* test. **(B)** Cep131 and PCM1 depletion disrupts centriolar protein localization to the centrosome. Left panels – confocal images of RPE-1 cells showing centriolar protein localization in PCM1 or Cep131, and double depleted cells with overexpressed Plk4. Cep192, Cep152, Plk4, and STIL, magenta and centrioles, green. Scale bar, 0.5 μm. Right panels – average normalized centrosome fluorescence intensities of centriolar proteins. Graph values expressed as means of biological replicates and SD. P value was determined using one-way ANOVA with Dunnett *post hoc* test. **(C)** Cep131-positive centriolar satellites are required for Unk localization to the centrosome. Left panels - confocal images of RPE-1 cells showing Unk localization in PCM1 or Cep131, and double depleted cells with overexpressed Plk4. Unk, magenta and centrioles, green. Scale bar, 5 μm. Insets scale bar, 1 μm. Middle panels – 8 μm radial fluorescence intensity and corresponding ratio quantification of Unk using Centrin as the centrosome centroid. Right panels - centrosomal Unk fluorescence intensity based on binned central 2 μm. Graph values expressed as the means of biological replicates and SD. P values were determined using one-way ANOVA with Dunnett post *hoc test*.

To elucidate how PCM1 and Cep131 differentially regulate Plk4-induced centriole overduplication, we quantified centrosome localization of the centriole assembly proteins Cep192, Cep152, Plk4, and STIL in PCM1, Cep131, or double knockdown (Fig. 4, B). PCM1 knockdown only reduced Plk4 at centrosomes, but did not affect centriole overduplication, suggesting that partial loss of Plk4 may not negatively affect centriole assembly or that other mechanisms compensate for this decrease in Plk4. Conversely, Cep131 knockdown reduced both Cep192 and Plk4 from centrosomes.

Cep131, knockdown decreased STIL at centrosomes by 40%, whereas PCM1 knockdown modestly increased centrosome localized STIL (Fig. 4, B). This suggests that both centriolar satellite proteins promote the localization of procentriole assembly proteins to the centrosome, but only Cep131 robustly promotes the localization of STIL. Plk4 phosphorylates STIL to recruit proteins necessary for centriole assembly^2^. The loss of STIL upon Cep131 depletion suggests this may attenuate centriole overduplication when Cep131 is lost. Because Unk knockdown reduces centrosomal Cep131 levels (Fig. 3, C), it would be expected that it also decreases STIL. However, we did not observe a significant change to STIL levels in Unk knockdown (Fig. 2, A) and suggest that, in contrast to the strong Cep131 knockdown, the residual levels of Cep131 in Unk knockdown are sufficient to localize STIL.

Because Cep131 and Unk knockdown both attenuate centriole overduplication, we asked if Cep131 or PCM1 regulate Unk localization. Cep131 knockdown reduced Unk levels at centrosomes (Fig. 4, C). Conversely, PCM1 knockdown increased Unk localization closer to the centrioles, although the overall centrosome levels of Unk were not affected (Fig. 4, C). These data suggest both Unk and Cep131 are centriolar satellite proteins that promote centriole overduplication. However, their mechanisms are different given their distinct effects on the localization of centriole assembly proteins.

### Unkempt’s RNA binding domain is required for Plk4-induced centriole overduplication

Unk’s RNA binding domain, composed of six zinc fingers in the N-terminus, is required for RNA binding and its control of cell morphology^40,41^. We asked if Unk RNA binding is required for Plk4-induced centriole overduplication. Cells were depleted of endogenous Unk and an siRNA-resistant (siR), N-terminal mCherry tag, Wild type (WTsiR) or RNA binding mutant (Mut.siR), were stably and inducibly expressed in S phase arrested cells, concurrent with Plk4 overexpression (Fig. 5A). Both WTsiR and Mut.siR proteins localized to centrosomes (Fig. S3, C). Depletion of endogenous Unk and expression of WTsiR, but not Mut.siR, with 0.4 µg/mL doxycycline rescued centriole overduplication. However, high expression of WTsiR with 1.0 µg/mL doxycycline, but not high expression of Mut.siR, blocked centriole overduplication (Fig. S3, D). In summary, these results suggest that 1) Unk expression levels affect Plk4-induced centriole overduplication and 2) Unk’s RNA binding domain is required for Plk4-induced centriole overduplication.

**Figure 5.**
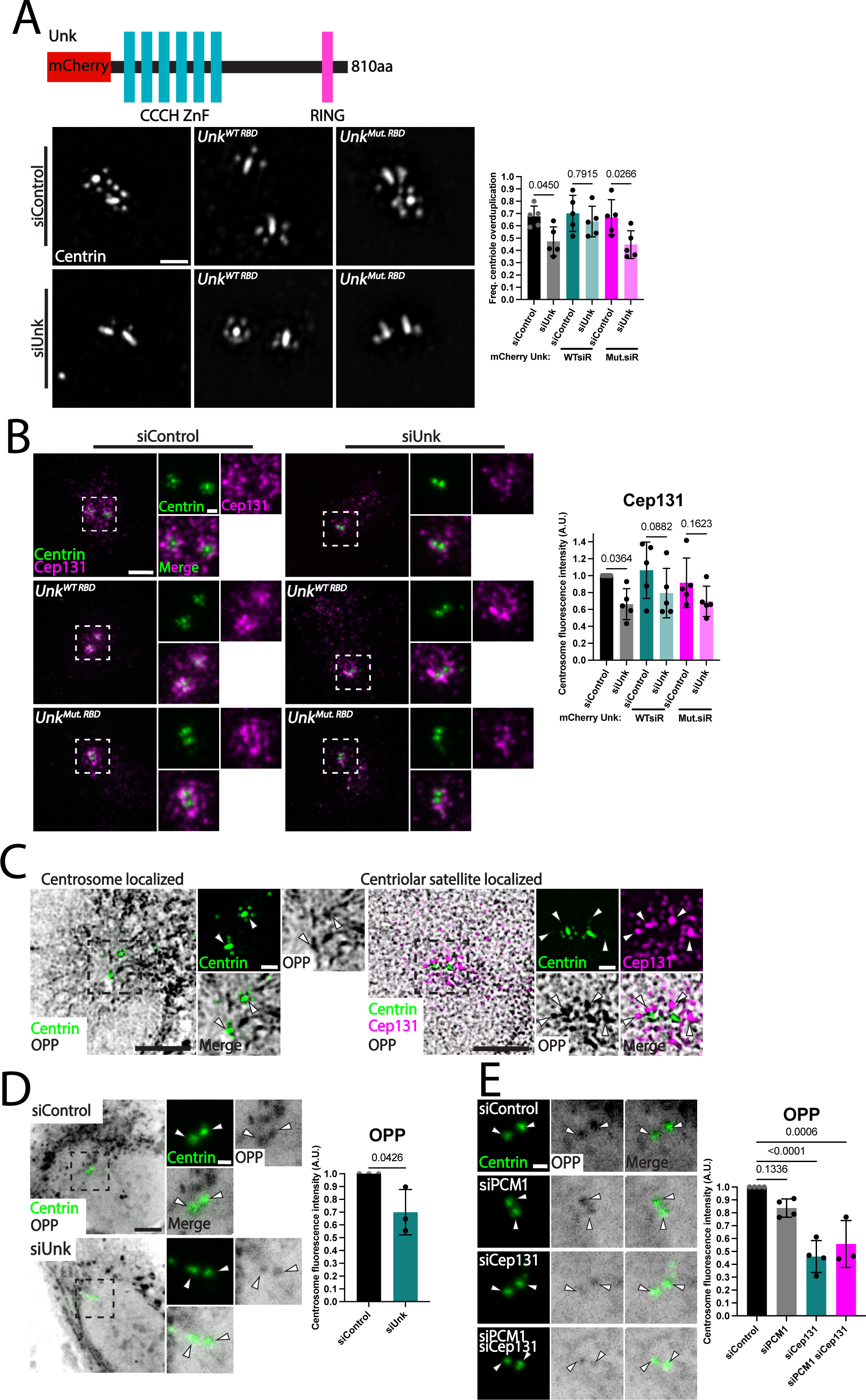
Unkempt’s RNA binding domain is required for Plk4-induced centriole overduplication. **(A)** Unk RNA binding is required for centriole overduplication. Left panels – amino acids mutated in Unk’s N terminal RNA binding domain to generate Mut.siR Unk. Middle panels – SIM images of RPE-1 cells showing centrioles in Unk depleted cells with overexpressed Plk4 and expression of either WTsiR or Mut.siR Unk. Centrioles, greyscale. Scale bar, 1 μm. Right panel – frequency of cells with centriole overduplication. Graph values expressed as the means of biological replicates and SD. P values were determined using one-way ANOVA with Šídák *post hoc* test. **(B)** Unk RNA binding is dispensable for Cep131-positive centriolar satellite localization. Left panels - confocal images of RPE-1 cells showing Cep131 localization in Unk depleted cells with overexpressed Plk4 and expression of either WTsiR or Mut.siR Unk. Cep131, magenta and centrioles, green. Scale bar, 5 μm. Insets scale bar, 1 μm. Right panels - centrosomal Cep131 fluorescence intensity based on binned central 2 μm. Graph values expressed as the means of biological replicates and SD. P values were determined using one-way ANOVA with Fishers Least Significant Difference (LSD) *post hoc* test. **(C)** Translation is enriched at centrosomes and centriolar satellites. Left panel - SIM showing centrosome localization of OPP in RPE-1 cells with overexpressed Plk4. OPP; greyscale and centrioles, green. Arrows indicate sites of OPP signal. Scale bar, 5 μm. Insets scale bar, 1 μm. Right panel – Confocal images showing centriolar satellite localization of OPP in RPE-1 cells with overexpressed Plk4. Cep131, magenta; OPP, greyscale; centrioles, green. Arrows indicate sites of OPP signal. Scale bar, 5 μm. Insets scale bar, 1 μm. **(D)** Unk suppresses translation at centrosomes during late centriole assembly. Left panels – SIM images of RPE-1 cells showing centrosome staining of OPP in Unk depleted cells 16 hours after Plk4 overexpression. OPP, greyscale and centrioles, green. Arrows indicate sites of OPP signal. Scale bar, 5 μm. Insets scale bar, 1 μm. Right panel - average normalized centrosome fluorescence intensities of OPP. Graph values expressed as the means of biological replicates and SD. P value was determined using an unpaired two-tailed t test. **(E)** Centriolar satellites promote local translation at the centrosome during late centriole assembly. Left panels – confocal images of RPE-1 cells showing centrosome staining of OPP in cells with PCM1 or Cep131, and double depleted cells 16 hours after Plk4 overexpression. OPP, greyscale and centrioles, green. Arrows indicate sites of OPP signal. Scale bar, 1 μm. Right panel - average normalized centrosome fluorescence intensities of OPP. Graph values expressed as the means of biological replicates and SD. P values were determined using one-way ANOVA with Dunnett *post hoc* test.

### Centriolar satellites and Unk promote local translation at and around centrosomes

Because Unk regulates mRNA translation^40,41,49^ and localizes to centrosomes and centriolar satellites, we investigated local translation at these structures. Cells were labeled with O-propargyl-puromycin (OPP). OPP is a puromycin analog that enters the ribosome acceptor site and is incorporated into nascent polypeptides. This modification can be fluorescently labeled to mark sites of translation within the cell^50^. RPE-1 cells were S phase arrested, induced for Plk4 overexpression for 16 hours, labeled with OPP, and co-stained with Cep131. OPP signal was observed throughout the cytoplasm and was enriched at and around centrosomes (Fig. 5, C and S4, B). OPP signal partially colocalized with Cep131 (Fig. 5, C). Moreover, OPP signal increased at centrosomes and decreased in the cytoplasm after 16 hours of PLK4 overexpression (Fig. S4, C). This suggests that centrosomes and Cep131-positive centriolar satellites surrounding centrosomes are responsive sites of active translation during S-phase. Though OPP labels truncated nascent polypeptides that are released from the ribosome, the possibility of diffusion or trafficking of these labeled nascent polypeptides to the centrosome cannot be eliminated^51^. Regardless, these results show that newly translated proteins localize to centrosomes and centriolar satellites in response to Plk4 overexpression and centriole overduplication. This is consistent with the mRNAs, RBPs, and translation machinery detected around centrosomes^29,31,52^.

Unk is an RNA binding protein previously shown to suppress translation^40,41,49^. Moreover, Unk localizes to centrosomes during early centriole assembly, and remains enriched at the centrosomes 6, 12, and 16 hours after Plk4 overexpression (Fig. 3, B, and 1, C). To determine whether Unk regulates translation at and around centrosomes, we depleted Unk and quantified OPP signal at centrosomes 16 hours after Plk4 overexpression. Unk depletion decreased centrosome translation by 35% without altering the levels of global translation (Fig. 5, D, Fig. S4, E).

Because Unk promotes the localization of centriolar satellites (Fig. 3, C) and centriolar satellites are sites of translation (Fig. 5, C), we asked whether centriolar satellites regulate local translation at centrosomes. To test this, PCM1, Cep131, or both were depleted and OPP signal at centrosomes was quantified 16 hours after Plk4 overexpression. Cep131 depletion decreased centrosome translation by 55% (Fig. 5, E). To explain the association of centriolar satellites with protein translation, Gene Ontology (GO) analysis was performed on published PCM1 and Cep131 BioID data (Fig. S4, D). The second most abundant GO category for protein regulatory functions interacting with PCM1 and Cep131 was RNA regulatory processes^25^. Unk and Cep131 centriolar satellite proteins interact with translation machinery including initiation and elongation factors, ribosomal proteins, and Poly A Binding Proteins (PABPs)^25^. These data suggest that Unk and centriolar satellites cooperate to promote local translation at centrosomes.

### Translation suppression occurs concomitantly with centriole overduplication

To determine if local translation is important for centriole overduplication, we monitored changes to centrosomal local translation at early timepoints after Plk4 overexpression. Cells were labeled with OPP at 6 and 12 hours after Plk4 overexpression, representing early stages of centriole overduplication (Fig. S2, D). Surprisingly, centrosome OPP signal decreased by approximately 45% and 30% at 6 and 12 hours, compared to the 0 hour timepoint (Fig. 6, A). This suggests that translation suppression occurs concomitantly with centriole overduplication. The decrease in centrosome translation during early centriole assembly events is in contrast to the late, 16 hour timepoint, where local translation is increased 40% (Fig. S4, C), highlighting a dynamic balance between suppressing and promoting local translation at centrosomes.

**Figure 6.**
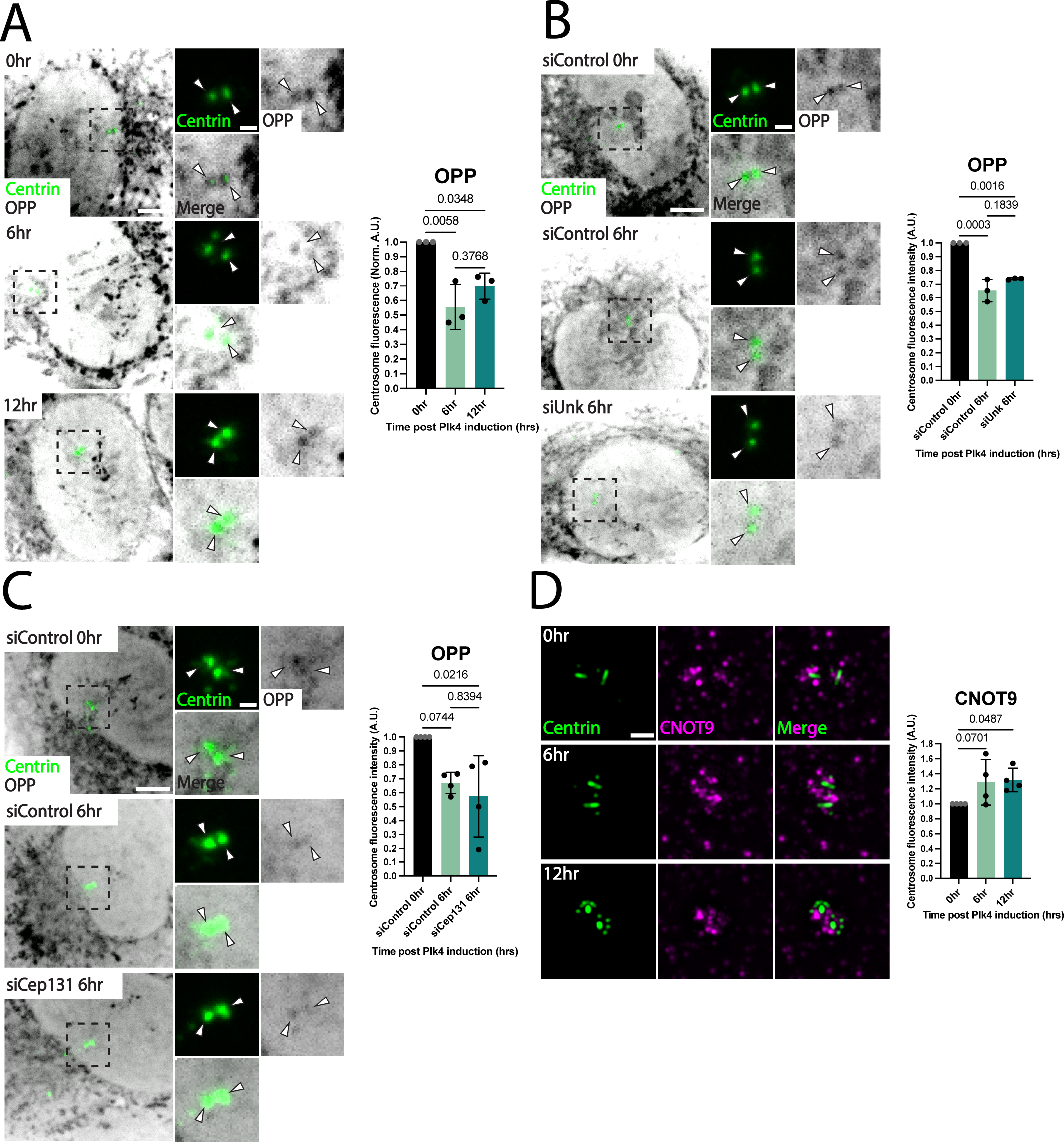
Translation suppression occurs concomitantly with centriole overduplication. **(A)** Plk4 overexpression causes reduced translation at centrosomes. Left panels – confocal images of RPE-1 cells showing centrosome staining of OPP 6 and 12 hours after Plk4 overexpression. OPP, greyscale and centrioles, green. Arrows indicate sites of OPP signal. Scale bar, 5 μm. Insets scale bar, 1 μm. Right panel – average normalized centrosome fluorescence intensities of OPP. Graph values expressed as the means of biological replicates and SD. P values were determined using one-way ANOVA with Dunnett *post hoc* test. **(B)** Unk does not regulate translation at the centrosome during early centriole assembly. Left panels – confocal images of RPE-1 cells showing centrosome staining of OPP in Unk depleted cells 6 hours after Plk4 overexpression. OPP, greyscale and centrioles, green. Arrows indicate sites of OPP signal. Scale bar, 5 μm. Insets scale bar,1 μm. Right panel - average normalized centrosome fluorescence intensities of OPP. Graph values expressed as the means of biological replicates and SD. P values were determined using one-way ANOVA with Šídák *post hoc* test. **(C)** Cep131 does not regulate translation at the centrosome during early centriole assembly. Left panels – confocal images of RPE-1 cells showing centrosome staining of OPP in Cep131 depleted cells 6 hours after Plk4 overexpression. OPP, greyscale and centrioles, green. Arrows indicate sites of OPP signal. Scale bar, 5 μm. Insets scale bar,1 μm. Right panel - average normalized centrosome fluorescence intensities of OPP. Graph values expressed as the means of biological replicates and SD. P values were determined using one-way ANOVA with Šídák *post hoc* test. **(D)** The CCR4-NOT complex subunit, CNOT9, localizes to the centrosome during early centriole assembly. Left panels – SIM images of RPE-1 cells showing centrosome staining of CNOT9 in cells 0, 6, and 12 hours after Plk4 overexpression. CNOT9, magenta and centrioles, green. Scale bar, 1 μm. Right panel - average normalized centrosome fluorescence intensities of CNOT9. Graph values expressed as the means of biological replicates and SD. P values were determined using one-way ANOVA with Fishers Least Significant Difference (LSD) *post hoc* test.

To elucidate whether Unk facilitates centriole overduplication by regulating translation at centrosomes during early centriole assembly, Unk depleted cells were stained for OPP 6 hours after Plk4 overexpression. Unk depletion did not change centrosomal OPP signal at this time (Fig. 6, B). This suggests that Unk does not regulate the reduction in local translation at early stages of Plk4-induced centriole assembly. Rather Unk affects translation later in the process when centrioles mature (Fig. 5, D).

To determine if centriolar satellites regulate local translation during early centriole assembly, Cep131 depleted cells were stained for OPP 6 hours after Plk4 overexpression. Like Unk depletion at this timepoint, Cep131 depletion did not change centrosomal OPP signal (Fig. 6, C). This suggests that neither Unk nor Cep131 regulate local translation during early centriole assembly. However, a balance of suppressing and promoting translation appears to be important during the dynamics of centriole assembly and maturation.

Unk is reported to suppress translation through its interactions with the RNA and translation suppressing complex, CCR4-NOT^49^. Specifically, Unk interacts with key effectors CNOT2 and CNOT9 to suppress translation^49^. Intriguingly, CNOT1, CNOT2, and CNOT3 are found in the centrosome proteome database^26^. To understand how Unk may be regulating translation suppression, members of these complexes were localized using immunofluorescence in Unk depleted cells, 16 hours after Plk4 overexpression. CNOT2 and CNOT9 both exhibited cytoplasmic and centrosome localization (Fig. S4, F). Unk knockdown reduced CNOT9 at centrosomes by 35%. In contrast, CNOT2 was not affected. Thus, Unk promotes the localization of the CNOT9 subunit of the CCR4-NOT translation suppression complex.

To understand how translation suppression is facilitated during early centriole overduplication, we asked whether the localization of CNOT9 changes 6 and 12 hours after Plk4 overexpression. Consistent with a role for CNOT9 in suppressing local translation at centrosomes, its levels increased at centrosomes by 28% and 31% 6 hours and 12 hours after Plk4 overexpression, respectively (Fig. 6, D). Therefore, the CCR4-NOT complex subunit CNOT9 increases in localization to centrosomes during early centriole assembly when translation is suppressed. These results suggest that CNOT9 is a regulator of the centrosome proximal translational program.

In summary, nascent protein translation decreases at and around centrosomes early (6- 12 hours) during Plk4-induced centriole duplication but then is elevated at late timepoints (16 hours). Consistent with the translation profile, the CCR4-NOT complex member, CNOT9, is elevated at early timepoints but then returns to normal by late timepoints (Fig. S4, G). This bimodal inactivation and activation of centrosome translation is controlled, in part, by Unk that promotes late localization of CNOT9. Because Unk promotes centrosome translation later in centriole assembly suggests Unk is not acting exclusively through CNOT9.

## Discussion

Centriole and centrosome amplification are commonly associated with cancers and can arise from dysregulation of the key regulatory kinase, Plk4. In this study, we discover Plk4 overexpression results in the increase of what, in essence, is a PLK4 cofactor for centriole assembly. Unk is an RNA binding protein that, in coordination with Cep131-positive centriolar satellites, promotes centriole overduplication and affects the regulation of local protein translation at centrosomes.

Centriolar satellites have both positive and negative effects on centriole duplication^20,53^. Here, the attenuation of centriole overduplication was observed upon Cep131, but not PCM1, knockdown. We favor a model in which centriole overduplication depends on centriolar satellite cargoes specifically scaffolded or regulated by Unk and Cep131. These cargoes may positively regulate centriole assembly or translationally suppress centriole assembly inactivators. Unk levels increase at centrosomes before, during, and remain increased after centrioles overduplicate. Cep131 levels at centrosomes remain static before, during, and decrease only after centrioles overduplicate. This suggests that centriolar satellites change throughout the duplication process, both in their composition and localization. This dynamic behavior is consistent with their cell cycle regulation and transport along MTs^24^. Moreover, Plk4 phosphorylates both PCM1 and Cep131 to maintain centriolar satellite integrity and stabilize Plk4 at the centrioles, respectively^22^. Perhaps centriolar satellites and Unk are required for efficient assembly at the start of centriole duplication in early S phase, but their role changes to control other centrosome regulatory events that are coordinated with the cell cycle, such as pericentriolar material accumulation and centriole maturation. In support of this hypothesis, proteins involved in both processes have been reported to interact with centriolar satellites^21,25,33,54^.

We found local translation occurs at centriolar satellites that cluster around centrosomes, near centrioles during S-phase. In support of this finding, translation machinery is reported to localize around centrosomes^55,56^, and proteomic studies identify association of centriolar satellites with translation machinery, as well as other RBPs^25,26,32^. In mouse multiciliated cells (MCC), OPP-positive, newly synthesized peptides colocalize with apical granules during centriole amplification. These granules are also positive for PCM1^57^. This supports the notion that centriolar satellites support local translation to promote centriole assembly. In the context of centriole overduplication, translation at the centrosome appears to depend on positive translation regulators like Unk and Cep131 during late centriole assembly, as well as negative translation regulators during early centriole assembly that may act through the CCR4-NOT complex. Suppressing translation of negative regulators of centriole duplication during early centriole duplication and promoting translation of positive regulators of centriole duplication by either Unk or Cep131 during late centriole duplication is one model by which centriolar satellites may facilitate centriole assembly. Depletion of Unk, Cep131, or PCM1 did not increase local translation, and this may be explained by spatial and temporal control of centriolar satellites and their specific translational targets that is beyond the resolution of this study^25,26^. For example, Unk may still suppress translation of certain mRNAs, consistent with its canonical role in translational regulation. OPP labeling does not identify which proteins are synthesized at the centrosome, so future work will be needed to identify these transcripts. In this model, Cep131-positive centriolar satellites localize translational machinery to centrosomes. Because Cep131 is also reduced from centrosomes when Unk is depleted, the observed phenotype is decreased translation at centrosomes during early and late centriole assembly. Interestingly, RBP-mediated translational suppression of mRNAs in neurons is required for their trafficking^58^. If translational suppression at centriolar satellites by Unk regulates centriolar satellite trafficking, then disruption of this site-specific suppression may affect centriolar satellite distribution, as is observed when Unk is depleted.

We show that the CCR4-NOT complex members, CNOT2 and CNOT9, localize to centrosomes and that CNOT9 increases in localization to centrosomes during early centriole assembly. The localization of this component of the translation suppression machinery at the centrosome correlates with translation suppression during early centriole assembly. CNOT9 may localize to centrioles during early centriole assembly to repress translation of centrosome proteins that would normally impair centriole assembly. In support of the centrosome-specific translation suppression model, knockdown of CNOT1 leads to translationally upregulated mRNAs that encode centrosome proteins^59^. Moreover, proximity labeling has identified CCR4- NOT complex members with centriolar proteins^60^. Requiring translation suppression for assembling and organizing macromolecular structures may be a conserved mechanism in cells. Indeed, the CNOT1 ortholog in *C. elegans*, LET-711, localizes to centrosomes and regulates spindle positioning and microtubule length^61^.

In summary, translation is regulated at centrosomes during Plk4-induced centriole overduplication and is mediated throughout the process by the RNA binding protein Unk and the centriolar satellite component Cep131, with Unk and Cep131 having roles in promoting centriole overduplication.

**Table 1.** Click Chemistry reagents.

| **Reagent** | **Final concentration** | **Catalog number** |
| --- | --- | --- |
| O-propargyl puromycin | 50 mM | Jena Bioscience NU-931-05 |
| Tris-Cl pH 7.0 | 10 mM |  |
| CuSO<sub>4</sub> | 100 mM |  |
| Cy5 picolyl azide | 6 mM | Vector Laboratories 1177-1 |
| THPTA | 2 mM |  |
| (+)-Sodium L-ascorbate | 1 mM | Sigma Life Science 1114-50G |

## Supplemental Figures

**Supplementary Figure 1 (A)** Plk4 overexpression promotes elevated UNK mRNA levels. Confocal images of RPE-1 cells showing UNK mRNA staining with endogenous and overexpressed Plk4. UNK mRNA, greyscale and centrioles, green. Scale bar, 5.0 μm. **(B)** siUnk #1 depletes whole cell levels of Unk protein in RPE-1 cells. Left – western blots of RPE-1 cell lysates with Unk knockdown using siRNA #1 with endogenous and overexpressed Plk4. Stained with PCNA as a loading control, and anti Unk. Right panels - fluorescence quantification of whole cell Unk protein levels with endogenous and overexpressed Plk4. Graph values expressed as means of biological replicates and SD. P values were determined using one-way ANOVA with Dunnett *post hoc* test. **(C)** Exogenous expression of WTsiR Unk localizes to centrosomes. Confocal images of RPE-1 cells showing expression of mCherry WT Unk at the centrosomes with Plk4 overexpression. Unk, magenta and centrioles, green. Scale bar, 5 μm. Insets scale bar, 1.0 μm. **(D)** siUnk #2 depletes whole cell levels of Unk protein in RPE-1 cells and attenuates centriole overduplication. Left panel – Confocal images of RPE-1 cells showing endogenous Unk protein in Unk depleted cells with Plk4 overexpression. Unk, magenta; Cep131, cyan; centrioles, green. Scale bar, 5.0 μm. Insets scale bar, 1.0 μm. Middle panel – frequency of cells with centriole overduplication. Graph values expressed as the means of biological replicates and SD. P value was determined using an unpaired two-tailed t test.

**Supplementary Figure 2 (A)** Knockdown of Unk does not affect total cellular protein levels of Plk4. Western blot analysis of RPE-1 cells depleted with siUnk #1. Cell lysates of RPE-1 cells with Unk knockdown using siRNA #1 with endogenous and overexpressed Plk4. Stained with PCNA as a loading control, and anti Plk4. **(B)** Unk depletion results in ectopic Centrin-positive foci that do not co-stain with Microtubules, PCM proteins, or centriole assembly proteins. Confocal images of RPE-1 cells showing Centrin-positive foci, green and stained for microtubules (DM1A), PCM proteins (PCNT and CDK5RAP2), and centriole assembly proteins (CEP152 and CEP192), magenta, in Unk depleted cells with endogenous and overexpressed Plk4. Scale bar, 5.0 μm. Arrows indicate sites of no colocalization with ectopic Centrin foci. **(C)** Unk depletion results in ectopic Centrin-positive foci that co-stain with centriolar satellite proteins. Confocal images of RPE-1 cells showing Centrin-positive foci and centriolar satellite protein localization in Unk depleted cells with endogenous and overexpressed Plk4. PCM1, Cep131, magenta and centrioles, green. Arrows indicate sites of colocalization with ectopic Centrin foci. Insets scale bar, 5.0 μm. **(D)** Centriole overduplication occurs in a time dependent manner after Plk4 overexpression. Frequency of RPE-1 cells with centriole overduplication 6 and 12 hours after Plk4 overexpression. Graph values expressed as the means of biological replicates and SD. P values were determined using one-way ANOVA with Dunnett *post hoc* test.

**Supplementary Figure 3 (A)** Centriolar satellites are reduced from the centrosome in Unk depleted cells with endogenous and overexpressed Plk4. Left panels – confocal images of RPE- 1 cells showing centriolar satellite localization in Unk depleted cells with endogenous and overexpressed Plk4. PCM1, Cep131, magenta and centrioles, green. Scale bar, 5 μm. Insets – SIM images of RPE-1 cells showing centrosome localization of centriolar satellites in Unk depleted cells with endogenous and overexpressed Plk4. PCM1, Cep131, magenta, and centrioles, green. Insets scale bar, 1.0 μm. Right, top panels – 8.0 μm radial fluorescence intensity of PCM1 and Cep131 using Centrin as the centrosome centroid. Right, middle panels - 8 μm ratio quantification of PCM1 and Cep131 using Centrin as the centroid. Right, bottom panels - centrosomal PCM1 and Cep131 fluorescence intensity based on binned central 2.0 μm. Graph values expressed as the means of replicates and SD. P values were determined using one-way ANOVA with Šídák *post hoc* test. **(B)** Knockdown of Unk does not affect total cellular protein levels of Cep131. Left panels - Western blot analysis of RPE-1 cells depleted with siUnk #1 with endogenous and overexpressed Plk4. Stained with Alpha tubulin, loading control, and anti Plk4. Right panels - fluorescence quantification of whole cell PCM1 and Cep131 protein levels with endogenous and overexpressed Plk4. Graph values expressed as means of biological replicates and SD. P values were determined using one-way ANOVA with Šídák *post hoc* test. **(C)** Exogenous expression of WTsiR and Mut.siR Unk localizes to centrosomes. Confocal images of RPE-1 cells showing expression of mCherry WTsiR and Mut.siR Unk at the centrosomes with Plk4 overexpression. Unk, magenta and centrioles, green. Scale bar, 5 μm. Insets scale bar, 1.0 μm. **(D)** High expression of WTsiR but not Mut.siR Unk blocks centriole overduplication. Frequency of cells with centriole overduplication. Graph values expressed as the mean of one biological replicate.

**Supplementary Figure 4 (A)** Unkempt’s RNA binding domain is dispensable for centriolar satellite localization. Left panels – 8.0 μm radial fluorescence intensity and corresponding ratio quantification of Cep131 using Centrin as the centrosome centroid. Right panels - centrosomal Cep131 fluorescence intensity based on binned central 2.0 μm. **(B)** OPP labels newly synthesized peptides. Confocal images of RPE-1 cells with controls for OPP labeling: OPP with click chemistry fluor reaction, click chemistry fluor reaction without OPP, OPP only, and DMSO only, all with Plk4 overexpression. OPP, magenta and centrioles, green. Scale bar, 5.0 μm **(C)** 16 hours of Plk4 overexpression causes reduced translation on a whole cell level, but increased translation at the centrosome. Left panel – fluorescence quantification of centrosomal OPP levels with endogenous and 16 hours Plk4 overexpression. Right panel - fluorescence quantification of whole cell OPP levels with endogenous and 16 hours of Plk4 overexpression. Graph values expressed as means of biological replicates and SD. P value was determined using an unpaired two-tailed t test**. (D)** Centriolar satellites interact with proteins involved in RNA regulation. Panther analysis of BioID data from PCM1 and Cep131 interacting proteins^25^ and their functional classification. **(E)** Unk does not impact translation on a whole cell level 16 hours of Plk4 overexpression. Left panel - fluorescence quantification of whole cell OPP levels with endogenous and 16 hours of overexpressed Plk4. Graph values expressed as the means of biological replicates and SD. P values were determined using an unpaired two-tailed t test. **(F)** Unk promotes the localization of CCR4-NOT translational suppression complex member, CNOT9. Left panels – confocal images of RPE-1 cells showing centrosome localization of CCR4-NOT complex proteins (CNOT9 and CNOT2) in Unk depleted cells with 16 hours of Plk4 overexpression. CNOT9, CNOT2, magenta and centrioles, green. Scale bar, 1 μm. Right panels – average normalized centrosome fluorescence intensities of CNOT9 and CNOT2. Graph values expressed as means of biological replicates and SD. P values were determined using an unpaired two-tailed t test. **(G)** The CCR4-NOT complex subunit, CNOT9, decreases at the centrosome 16 hours of Plk4 overexpression. Average normalized centrosome fluorescence intensities of CNOT9. Graph values expressed as two means of biological replicates and the range. P values were determined using an unpaired two-tailed t test.

## Materials and methods

### Cell culture growth conditions and small interfering RNA treatments

RPE-1-Tet-Plk4 Cetn2-GFP (kind gift of M.B. Tsou)^39^ were grown in DMEM/F12 with glutamine (Cytiva or Life Technologies) with 10% tetracycline-free fetal bovine serum (FBS; Peak Serum) supplemented with penicillin and streptavidin (Life Technologies) at 37°C with 5% CO2. A PCR test for mycoplasma contamination was performed every 6 months. For Plk4 overexpression experiments, cells were plated onto 12-mm circular cover glass, No. 1.5 (Electron Microscopy Sciences) coated with collagen (Sigma, C9791) in 24-well plates at 12,500 cells and 7,000 cells for 48- and 72-hour knockdown, respectively. The following day, cells were treated with small interfering RNAs using the Lipofectamine RNAiMAX Transfection Reagent (Thermo Fisher Scientific) following the manufacturer’s instructions. Transfection complexes were removed from cells 5 - 6 hours after addition, and cells were provided fresh medium. 24 hours prior to fixation, cells were arrested with 1.5 µg/mL aphidicolin (Cayman Chemical Company). Plk4 was induced with 1.0 µg/mL doxycycline (Sigma-Aldrich) 8 hours after initiation of the aphidicolin arrest.

### Generation of Unk RNAi resistant constructs

To generate WTsiR and Mut.siR Unk constructs, 1.8 kb gene blocks (Integrated DNA Technologies) encompassing the N terminal region of Unk, six Zinc fingers, and siRNA target regions were synthesized. Zinc finger residues, previously shown to abrogate Unk RNA binding when mutated to alanine^41^ (R119A, Y120A, N143A, F149A, Q228A, F289A, R310A, F316A), were utilized for Mut.siR Unk. For generating RNAi resistance towards two different siRNAs, synonymous codon optimized mutations were made to siRNA #1 (Thermo fisher HSS150335); GTGCCCTCCTCTGTAGAAACAGCAG, siRNA #2 (pool of 4 siRNAs) (Horizon Discovery siGENOME M-022950-01-0010); CCTGAAAGAATTCCGCACA, AAGCACAAATACAGGTCGT, CCACCAAGTGCAACGACAT, GGAGAAGACTTTCGATAAC, targets. To generate WTsiR and Mut.siR Unk, Human Unk was first amplified from cDNA (Origene SC315847) using Fwd: 5’ ACAGTCGACATGTCGAAGGGCCCCGG 3’ and Rev: 5’ TCACCGGTTCACGACTGGAGGGTGTGG 3’ and cloned into pBluescript II KS (-) using SalI and AgeI restriction enzyme sites. EcoRI and HindIII sites were appended to the ends of WTsiR and Mut.siR gene blocks, digested, and cloned into the pBluescript II KS (-) with full length Unk. Full length WTsiR and Mut.siR Unk from pBluescript II KS (-) was amplified and cloned into the lentivirus pCW57.1 N-term mCherry-BLAST construct for inducible doxycycline expression of pCW57.1-mCherry-WTsiR Unk and pCW57.1-mCherry-Mut.siR Unk. Sequencing of these constructs was confirmed using nanopore sequencing.

### Lentivirus transduction

Plasmids were isolated using ZymoPURE Plasmid Miniprep (D4208T). HEK293T cells at 60% confluency in 10 cm dishes were transfected with 1.5 μg of psPAX2 second generation lentiviral packaging plasmid (Addenge 12260), 0.5 mg of pMD2.G envelope expressing plasmid, and 2 mg of pCW57-mCherry-WTsiRUnk and pCW57-mCherry-Mut.siRUnk in Opti-MEM (Life Technologies 31985070), using the Lipofectamine 2000 Transfection Reagent (Thermo Fisher 11668027) according to the manufacturer’s instructions. The following two days, medium containing viral particles was removed from HEK293T cells and added to RPE-1/Tet-Plk4 Cetn2-GFP with 10 µg/mL of Polybrene. Transduced cells were selected with 10 µg/mL Blasticidin for one week prior to isolating clonal cell lines.

### Immunofluorescence

Cells were fixed in cold MeOH for 8 minutes, washed three times in phosphate-buffered saline (PBS), and blocked in Knudsen buffer (1× PBS, 0.5% bovine serum albumin, 0.5% NP-40, 1.0 mM MgCl2, 1.0 mM NaN3) for 1 hour. Cells stained for tubulin were fixed with glutaraldehyde and formaldehyde, as previously described^62^. Antibody staining was conducted at room temperature for 1 - 2 hours. Samples were washed 3 times for 5 minutes each in PBS prior to secondary antibody and DNA staining in Knudson buffer for 1 hour, except for samples stained for tubulin in which fix, staining, and wash were conducted in PHEM buffer. Primary antibodies: 1:800 rabbit α-Unk (Sigma HPA023636); 1:1000 rabbit α-STIL (the generous gift of J. Reiter, Department of Biochemistry and Biophysics, University of California, San Francisco School of Medicine, San Francisco, CA); rabbit α-CDK5RAP2 (Bethyl 50-157-1418); 1:1000 rabbit α-Plk4 (the generous gift of A. Holland, Department of Molecular Biology and Genetics, Johns Hopkins University School of Medicine, Baltimore, MD); 1:5,000 Guinea Pig α-CEP131 (the generous gift of J. Reiter), Department of Biochemistry and Biophysics, University of California, San Francisco School of Medicine, San Francisco, CA); 1:2500 rabbit α-CEP152 (Bethyl A302- 480A); rabbit 1:2000 α-CEP192 (the generous gift of A. Holland, Department of Molecular Biology and Genetics, Johns Hopkins University School of Medicine, Baltimore, MD); rabbit 1:2000 α-PCM1, 1:1000 α-Cep63 (Proteintech 16268-1-AP); (Bethyl A301-150A); 1:2000 rabbit α-PCNT (Abcam ab4448); 1:300 mouse α-tubulin (Sigma CP06-100UG); 1:100 rabbit α-CNOT2 (Proteintech 10313-1-AP); 1:100 rabbit α-CNOT9 (Proteintech 22503-1-AP). Secondary staining: 1:1000 Alexa anti-rabbit 594, Alexa anti-mouse 594, and Alexa anti-guinea pig 647, respectively (Thermo Fisher Scientific). Where appropriate, DNA was stained using Hoechst 33342 (Thermo Fisher Scientific 62249). Coverslips were mounted using Citifluor (Ted Pella) and sealed with clear nail polish for Confocal imaging and ProLong Gold Antifade (Thermo Fisher Scientific P10144) for SIM.

### Fluorescence imaging

Images were collected using a Yokogawa X1 spinning disk confocal (SDC) on a Nikon Ti-E inverted microscope stand with 100X Plan Apo NA 1.4 objective. Images were acquired at room temperature using 0.25 μm Z steps on an Andor iXon EM-CCD camera with exposure settings between 0-500 ms and 0-150 intensification, depending upon the experiment, and no binning of pixels. SIM images were acquired using a Nikon SIM (N-SIM) on a Nikon Ti2 (Nikon Instruments; LU-N3-SIM) microscope equipped with a 100× SR Apo TIRF, NA 1.49 objective. Images were captured using a Hamamatsu ORCA-Flash 4.0 Digital CMOS camera (C13440) with 0.1-0.2μm Z step sizes. Exposure settings were between 0-300 ms depending upon the experiment. All images were collected at 25°C using NIS Elements software (Nikon). Raw SIM images were reconstructed using the image slice reconstruction algorithm (NIS Elements).

### Centriole counts

GFP-Centrin fluorescence was utilized to count centrioles manually using the 100× PlanApo DIC, NA 1.4 objective with a 1.5X magnification optivar on a Nikon TiE inverted microscope stand. Fields of view were randomly chosen.

### Image analysis

Image analysis was performed using FIJI^63^. Image stacks were projected by maximum intensity. Fluorescence intensities of centrosomes were measured within a 1-μm radius encompassing each individual centrosome. A measurement of cell intensity near each centrosome was acquired for non-centrosome signal background subtraction. To measure the fluorescence intensity of centriolar satellite proteins, fluorescence was measured within an 8.0-μm radius encompassing both centrosomes and the surrounding cytoplasm. The lowest intensity value found in the 8.0-μm radius was subtracted from the calculated radial intensities. If subtraction of background signal generated a negative fluorescence signal, it was converted to 0. For radial fluorescence intensity analyses, we utilized the Radial Profile Extended algorithm (ImageJ). Briefly, maximum intensity-projected images were selected for analysis. Centrosomes and cells identified by the algorithm were manually confirmed for accuracy prior to the algorithm calculating the in-cell radial fluorescence intensity.

To assess total fluorescence within a cell, background was subtracted from an average 5 x 5 μm box outside of the cell. The Centrin channel was thresholded to generate a binary image encompassing the cell boundaries. This was subjected to the erode, dilate, and fill holes binary functions followed by applying the create selection function. The selection was added to the region of interest manager and then transferred to the channel to be quantified.

### Statistical methods and data collection

Centriole counts for two compared groups were analyzed using a t test with two tails. Fluorescence intensity measurements were compared using an unpaired t test with two tails. All statistics were calculated using Prism (Graphpad). For multiple, or more than two comparisons, a One-Way ANOVA with *post hoc* tests of Dunnett, Šidák, or Fisher’s Least Significant Difference (LSD) were conducted where indicated. Investigators were not blinded when collecting data. Images were collected identically within experiments and data analysis was automated to the extent possible to prevent bias.

### Single molecule inexpensive Fluorescence in situ Hybridization (smiFISH)

Probes were generated using Stellaris probe design for the *Unkempt* gene and set to generate 48 probes. Probes were ordered from Integrated DNA Technologies (IDT). The reverse complement of the X FLAP sequence; CCTCCTAAGTTTCGAGCTGGACTCAGTG, was added to the 5’ end of each probe. The 48 probes were resuspended to a final concentration of 100 μM and then combined to make a final equimolar probe mix of 100 μM. To create smiFISH duplexes, a 10 μL mixture consisting of a final concentration of 20 μM equimolar probe mix, 1x NEB3 buffer, and 25 μM Alexa Flour 647 (IDT) was combined. The duplexes were then assembled in a thermocycler with the following parameters: 85°C for 3 minutes, 65°C for 3 minutes, 25°C for 5 minutes, and then kept on ice or frozen at −20°C for storage. For RNA hybridization, cells were fixed onto 12 mm coverslips according to the LGC Biosearch Technologies RNA FISH protocol. For each 12 mm coverslip, 2 μL of smiFISH duplexes in 50 μL of Stellaris RNA FISH Hybridization buffer (LGC Biosearch Technologies SMF-HB1-10) was incubated on coverslips overnight at 37°C and washed with Stellaris RNA FISH Wash Buffer A (LGC Biosearch Technologies SMF-WA1-60) 3 times for 5 minutes, followed by one final wash in Stellaris RNA Wash Buffer B (LGC Biosearch Technologies SMF-WB1-20).

### *O-propargyl puromycin labeling and* click chemistry *on fixed cells*

Cells were labeled with O-propargyl puromycin (OPP) (Jena Bioscience NU-931-05) at a final concentration of 50 μM in pre-warmed media for 15 minutes. Cells were then fixed in methanol for 8 minutes. Following fixation and 3 PBS washes, 500 μL total of click chemistry reagents at the final concentration of 10 mM Tris-Cl pH 7.0, 100 mM CuSO4, 6 mM Cy5 picolyl azide (Vector Laboratories 1177-1), 2 mM THPTA, 1 mM freshly made (+)-Sodium L-ascorbate (Sigma Life Science 1114-50G), and DEPC-treated water were added per coverslip in a 24 well plate for 30 minutes at room temperature protected from light. Coverslips were washed 3 times with PBS for 5 minutes and either mounted or processed for antibody staining. For antibody staining, cells were permeabilized, blocked for 1 hour, and incubated in primary antibody for 1 hour followed by 3, 5-minute washes in PBS. Secondary antibodies were incubated for 1 hour, followed by 3, 5-minute washes in PBS. Cells were imaged on the same day. A decrease in OPP signal was observed following primary and secondary incubation overnight.

### Western blot

For western blot, cells were lysed with whole cell lysis buffer (20 mM HEPES-KOH pH 7.9, 10% glycerol, 300 mM KCl, 0.1% IGEPAL, 1 mM DTT and 1X protease inhibitor cocktail (Thermo scientific PI78440); 100 μL of lysis buffer per 1×10^6^cells) by gentle inversion at 4°C for 30 minutes. After lysis the tubes were spun at 17,000 x g for 30 minutes and the supernatants were collected. The supernatants were mixed with an equal volume of lysis buffer without salt producing a final salt concentration to 150 mM KCl. Total protein was quantified using the Quick Start Bradford Protein Assay (Bio-Rad 5000201). Approximately 30 μg of protein was mixed with 1x SDS loading buffer and boiled for 10 minutes. Western blots were blocked with TBS + Tween-20 at 0.05% and 0.5% BSA for one hour at room temperature and incubated in primary overnight in TBST+0.5% BSA. After 3 TBST washes, blots were incubated in Licor IR680 or IR800 secondary antibodies at 1:20,000 for 1 hour, washed 3 times with TBST and imaged using a fluorescence Odyssey CLx Imager and LI-COR Acquisition Software (LICORbio).

### RNA sequencing

RNA used for polyA-selected library generation for mRNA sequencing was isolated with the Monarch Total RNA isolation miniprep kit (New England Biolabs). Libraries were constructed using the Nugen Universal Plus mRNA-SEQ library construction kit (Nugen 0508) and sequenced on an Illumina NovaSEQ 6000 sequencer by the Genomics and Microarray shared resource at the University of Colorado Cancer Center.

### Bioinformatics

Bioinformatics was performed as previously described in ^45^. Briefly, RNA-seq libraries were sequenced using an Illumina NovaSeq 6000 (2 × 150) to a depth of 60–100 million paired-end reads. Reads were trimmed using cutadapt (v1.16) to remove adapters and aligned to the hg38 genome using STAR v2.5.2a^64^. Gene counts were quantified using feature Counts v1.6.2^65^. Genes that were differentially regulated between endogenous and Plk4 overexpression were analyzed using DESeq2 v1.28.1^66^.

## Acknowledgments

We are grateful to present and past Pearson lab members who provided insight and comments on the project. Jernej Murn provided reagents and helpful advice, Marisa Ruehle assisted with OPP and smFISH experimental methods, Andrew Holland and Jeremy Reiter generously shared antibodies, and Meng-Fu Bryan Tsou supplied the RPE1-Tet-Plk4/GFP-Cetn-2 cell line. This research was funded by NIH-NIGMS R35 GM140813 (CGP), R35 GM140813 Diversity Supplement (AM), R35 GM133385 (JMT) and W.M. Keck Foundation (JMT and CGP).

## Author contributions

AM, ASW, and CGP designed research project. AM performed most of the experiments, analyzed, and interpreted data under the guidance of CGP. RNA sequencing analysis was performed by RMS. All the authors participated in writing the manuscript, approved the final version to be published, and agreed to be accountable for all aspects of this study.

The authors declare no competing financial interests.

## Abbreviations

RBP: RNA binding protein
Unk: Unkempt
MTs: Microtubules
CA: Centrosome amplification

## References

1. Naslavsky N, Caplan S. Endocytic membrane trafficking in the control of centrosome function. Curr Opin Cell Biol. Aug 2020;65:150–155. doi:10.1016/j.ceb.2020.01.009

2. Moyer TC, Holland AJ. PLK4 promotes centriole duplication by phosphorylating STIL to link the procentriole cartwheel to the microtubule wall. Elife. May 22 2019;8doi:10.7554/eLife.46054

3. Kleylein-Sohn J, Westendorf J, Le Clech M, Habedanck R, Stierhof YD, Nigg EA. Plk4-induced centriole biogenesis in human cells. Dev Cell. Aug 2007;13(2):190–202. doi:10.1016/j.devcel.2007.07.002

4. Nigg EA, Holland AJ. Once and only once: mechanisms of centriole duplication and their deregulation in disease. Nat Rev Mol Cell Biol. May 2018;19(5):297–312. doi:10.1038/nrm.2017.127

5. Coelho PA, Bury L, Shahbazi MN, et al. Over-expression of Plk4 induces centrosome amplification, loss of primary cilia and associated tissue hyperplasia in the mouse. Open Biol. Dec 2015;5(12):150209. doi:10.1098/rsob.150209

6. Denu RA, Shabbir M, Nihal M, et al. Centriole Overduplication is the Predominant Mechanism Leading to Centrosome Amplification in Melanoma. Mol Cancer Res. Mar 2018;16(3):517–527. doi:10.1158/1541-7786.Mcr-17-0197

7. Ganapathi Sankaran D, Stemm-Wolf AJ, Pearson CG. CEP135 isoform dysregulation promotes centrosome amplification in breast cancer cells. Mol Biol Cell. May 1 2019;30(10):1230–1244. doi:10.1091/mbc.E18-10-0674

8. Zhou H, Kuang J, Zhong L, et al. Tumour amplified kinase STK15/BTAK induces centrosome amplification, aneuploidy and transformation. Nature genetics. 1998;20(2):189–193.

9. Pihan GA, Purohit A, Wallace J, et al. Centrosome defects and genetic instability in malignant tumors. Cancer Res. Sep 1 1998;58(17):3974–85.

10. Ghadimi BM, Sackett DL, Difilippantonio MJ, et al. Centrosome amplification and instability occurs exclusively in aneuploid, but not in diploid colorectal cancer cell lines, and correlates with numerical chromosomal aberrations. Genes Chromosomes Cancer. Feb 2000;27(2):183–90.

11. Marthiens V, Rujano MA, Pennetier C, Tessier S, Paul-Gilloteaux P, Basto R. Centrosome amplification causes microcephaly. Nature cell biology. 2013;15(7):731–740.

12. Levine MS, Bakker B, Boeckx B, et al. Centrosome Amplification Is Sufficient to Promote Spontaneous Tumorigenesis in Mammals. Developmental Cell. 2017/02/06/ 2017;40(3):313-322.e5. 10.1016/j.devcel.2016.12.022

13. Brown NJ, Marjanović M, Lüders J, Stracker TH, Costanzo V. Cep63 and cep152 cooperate to ensure centriole duplication. PLoS One. 2013;8(7):e69986. doi:10.1371/journal.pone.0069986

14. Sonnen KF, Gabryjonczyk AM, Anselm E, Stierhof YD, Nigg EA. Human Cep192 and Cep152 cooperate in Plk4 recruitment and centriole duplication. J Cell Sci. Jul 15 2013;126(Pt 14):3223–33. doi:10.1242/jcs.129502

15. Ohta M, Ashikawa T, Nozaki Y, et al. Direct interaction of Plk4 with STIL ensures formation of a single procentriole per parental centriole. Nature Communications. 2014/10/24 2014;5(1):5267. doi:10.1038/ncomms6267

16. Čajánek L, Glatter T, Nigg EA. The E3 ubiquitin ligase Mib1 regulates Plk4 and centriole biogenesis. J Cell Sci. May 1 2015;128(9):1674–82. doi:10.1242/jcs.166496

17. Cunha-Ferreira I, Bento I, Pimenta-Marques A, et al. Regulation of autophosphorylation controls PLK4 self-destruction and centriole number. Curr Biol. Nov 18 2013;23(22):2245–2254. doi:10.1016/j.cub.2013.09.037

18. Rogers GC, Rusan NM, Roberts DM, Peifer M, Rogers SL. The SCF Slimb ubiquitin ligase regulates Plk4/Sak levels to block centriole reduplication. J Cell Biol. Jan 26 2009;184(2):225–39. doi:10.1083/jcb.200808049

19. Kim DH, Ahn JS, Han HJ, et al. Cep131 overexpression promotes centrosome amplification and colon cancer progression by regulating Plk4 stability. Cell Death Dis. Jul 29 2019;10(8):570. doi:10.1038/s41419-019-1778-8

20. Stemm-Wolf AJ, O’Toole ET, Sheridan RM, Morgan JT, Pearson CG. The SON RNA splicing factor is required for intracellular trafficking structures that promote centriole assembly and ciliogenesis. Molecular Biology of the Cell. 2021;32(20):ar4. doi:10.1091/mbc.E21-06-0305

21. Conkar D, Bayraktar H, Firat-Karalar EN. Centrosomal and ciliary targeting of CCDC66 requires cooperative action of centriolar satellites, microtubules and molecular motors. Sci Rep. Oct 3 2019;9(1):14250. doi:10.1038/s41598-019-50530-4

22. Staples CJ, Myers KN, Beveridge RD, et al. The centriolar satellite protein Cep131 is important for genome stability. J Cell Sci. Oct 15 2012;125(Pt 20):4770–9. doi:10.1242/jcs.104059

23. Aydin ÖZ, Taflan SO, Gurkaslar C, Firat-Karalar EN. Acute inhibition of centriolar satellite function and positioning reveals their functions at the primary cilium. PLoS biology. 2020;18(6):e3000679.

24. Kubo A, Sasaki H, Yuba-Kubo A, Tsukita S, Shiina N. Centriolar satellites: molecular characterization, ATP-dependent movement toward centrioles and possible involvement in ciliogenesis. J Cell Biol. Nov 29 1999;147(5):969–80. doi:10.1083/jcb.147.5.969

25. Gheiratmand L, Coyaud E, Gupta GD, et al. Spatial and proteomic profiling reveals centrosome-independent features of centriolar satellites. Embo j. Jul 15 2019;38(14):e101109. doi:10.15252/embj.2018101109

26. Gupta GD, Coyaud É, Gonçalves J, et al. A Dynamic Protein Interaction Landscape of the Human Centrosome-Cilium Interface. Cell. Dec 3 2015;163(6):1484–99. doi:10.1016/j.cell.2015.10.065

27. Lerit DA. Signed, sealed, and delivered: RNA localization and translation at centrosomes. Mol Biol Cell. May 1 2022;33(5)doi:10.1091/mbc.E21-03-0128

28. Safieddine A, Coleno E, Salloum S, et al. A choreography of centrosomal mRNAs reveals a conserved localization mechanism involving active polysome transport. Nature Communications. 2021/03/01 2021;12(1):1352. doi:10.1038/s41467-021-21585-7

29. Filippova N, Yang X, King P, Nabors LB. Phosphoregulation of the RNA-binding protein Hu antigen R (HuR) by Cdk5 affects centrosome function. J Biol Chem. Sep 14 2012;287(38):32277–87. doi:10.1074/jbc.M112.353912

30. Jao LE, Akef A, Wente SR. A role for Gle1, a regulator of DEAD-box RNA helicases, at centrosomes and basal bodies. Mol Biol Cell. Jan 1 2017;28(1):120–127. doi:10.1091/mbc.E16-09-0675

31. Ishigaki Y, Nakamura Y, Tatsuno T, Hashimoto M, Iwabuchi K, Tomosugi N. RNA-binding protein RBM8A (Y14) and MAGOH localize to centrosome in human A549 cells. Histochemistry and cell biology. 2014;141:101–109.

32. Arslanhan MD, Gulensoy D, Firat-Karalar EN. A Proximity Mapping Journey into the Biology of the Mammalian Centrosome/Cilium Complex. Cells. 2020;9(6):1390.

33. Dammermann A, Merdes A. Assembly of centrosomal proteins and microtubule organization depends on PCM-1. J Cell Biol. Oct 28 2002;159(2):255–66. doi:10.1083/jcb.200204023

34. Garapaty S, Mahajan MA, Samuels HH. Components of the CCR4-NOT complex function as nuclear hormone receptor coactivators via association with the NRC-interacting Factor NIF-1. J Biol Chem. Mar 14 2008;283(11):6806–16. doi:10.1074/jbc.M706986200

35. Singh CK, Denu RA, Nihal M, et al. PLK4 is upregulated in prostate cancer and its inhibition reduces centrosome amplification and causes senescence. The Prostate. 2022;82(9):957–969. 10.1002/pros.24342

36. Peel N, Stevens NR, Basto R, Raff JW. Overexpressing centriole-replication proteins in vivo induces centriole overduplication and de novo formation. Current Biology. 2007;17(10):834–843.

37. Bettencourt-Dias M, Rodrigues-Martins A, Carpenter L, et al. SAK/PLK4 is required for centriole duplication and flagella development. Current biology. 2005;15(24):2199–2207.

38. Habedanck R, Stierhof Y-D, Wilkinson CJ, Nigg EA. The Polo kinase Plk4 functions in centriole duplication. Nature cell biology. 2005;7(11):1140–1146.

39. Kim M, O’Rourke BP, Soni RK, Jallepalli PV, Hendrickson RC, Tsou M-FB. Promotion and suppression of centriole duplication are catalytically coupled through PLK4 to ensure centriole homeostasis. Cell reports. 2016;16(5):1195–1203.

40. Murn J, Zarnack K, Yang YJ, et al. Control of a neuronal morphology program by an RNA-binding zinc finger protein, Unkempt. Genes Dev. Mar 1 2015;29(5):501–12. doi:10.1101/gad.258483.115

41. Murn J, Teplova M, Zarnack K, Shi Y, Patel DJ. Recognition of distinct RNA motifs by the clustered CCCH zinc fingers of neuronal protein Unkempt. Nat Struct Mol Biol. Jan 2016;23(1):16–23. doi:10.1038/nsmb.3140

42. Lambrus BG, Uetake Y, Clutario KM, et al. p53 protects against genome instability following centriole duplication failure. J Cell Biol. Jul 6 2015;210(1):63–77. doi:10.1083/jcb.201502089

43. Kodani A, Yu TW, Johnson JR, et al. Centriolar satellites assemble centrosomal microcephaly proteins to recruit CDK2 and promote centriole duplication. Elife. Aug 22 2015;4doi:10.7554/eLife.07519

44. Hall EA, Keighren M, Ford MJ, et al. Acute versus chronic loss of mammalian Azi1/Cep131 results in distinct ciliary phenotypes. PLoS genetics. 2013;9(12):e1003928.

45. Stemm-Wolf AJ, O’Toole ET, Sheridan RM, Morgan JT, Pearson CG. The SON RNA splicing factor is required for intracellular trafficking structures that promote centriole assembly and ciliogenesis. Mol Biol Cell. Oct 1 2021;32(20):ar4. doi:10.1091/mbc.E21-06-0305

46. Jewett CE, McCurdy BL, O’Toole ET, et al. Trisomy 21 induces pericentrosomal crowding delaying primary ciliogenesis and mouse cerebellar development. Elife. Jan 19 2023;12doi:10.7554/eLife.78202

47. Hall EA, Kumar D, Prosser SL, et al. Centriolar satellites expedite mother centriole remodeling to promote ciliogenesis. Elife. 2023;12:e79299.

48. Hall EA, Kumar D, Prosser SL, et al. Centriolar satellites expedite mother centriole remodeling to promote ciliogenesis. Elife. Feb 15 2023;12doi:10.7554/eLife.79299

49. Shah K, He S, Turner DJ, et al. Regulation by the RNA-binding protein Unkempt at its effector interface. Nat Commun. Apr 11 2024;15(1):3159. doi:10.1038/s41467-024-47449-4

50. Staudacher J, Rebnegger C, Gasser B. Treatment with surfactants enables quantification of translational activity by O-propargyl-puromycin labelling in yeast. BMC Microbiology. 2021/04/20 2021;21(1):120. doi:10.1186/s12866-021-02185-3

51. Enam SU, Zinshteyn B, Goldman DH, et al. Puromycin reactivity does not accurately localize translation at the subcellular level. Elife. Aug 26 2020;9doi:10.7554/eLife.60303

52. Pascual R, Segura-Morales C, Omerzu M, et al. mRNA spindle localization and mitotic translational regulation by CPEB1 and CPEB4. RNA. 2021;27(3):291–302.

53. Aydin Ö Z, Taflan SO, Gurkaslar C, Firat-Karalar EN. Acute inhibition of centriolar satellite function and positioning reveals their functions at the primary cilium. PLoS Biol. Jun 2020;18(6):e3000679. doi:10.1371/journal.pbio.3000679

54. Conkar D, Culfa E, Odabasi E, Rauniyar N, Yates JR, 3rd, Firat-Karalar EN. The centriolar satellite protein CCDC66 interacts with CEP290 and functions in cilium formation and trafficking. J Cell Sci. Apr 15 2017;130(8):1450–1462. doi:10.1242/jcs.196832

55. Kwon OS, Mishra R, Safieddine A, et al. Exon junction complex dependent mRNA localization is linked to centrosome organization during ciliogenesis. Nature communications. 2021;12(1):1351.

56. Sepulveda G, Antkowiak M, Brust-Mascher I, et al. Co-translational protein targeting facilitates centrosomal recruitment of PCNT during centrosome maturation in vertebrates. Elife. Apr 30 2018;7doi:10.7554/eLife.34959

57. Liu H, Li H, Jiang Z, et al. A local translation program regulates centriole amplification in the airway epithelium. Scientific Reports. 2023;13(1):7090.

58. Das S, Vera M, Gandin V, Singer RH, Tutucci E. Intracellular mRNA transport and localized translation. Nature Reviews Molecular Cell Biology. 2021;22(7):483–504.

59. Gillen SL, Giacomelli C, Hodge K, Zanivan S, Bushell M, Wilczynska A. Differential regulation of mRNA fate by the human Ccr4-Not complex is driven by coding sequence composition and mRNA localization. Genome Biol. Oct 6 2021;22(1):284. doi:10.1186/s13059-021-02494-w

60. Youn J-Y, Dunham WH, Hong SJ, et al. High-density proximity mapping reveals the subcellular organization of mRNA-associated granules and bodies. Molecular cell. 2018;69(3):517–532. e11.

61. DeBella LR, Hayashi A, Rose LS. LET-711, the Caenorhabditis elegans NOT1 ortholog, is required for spindle positioning and regulation of microtubule length in embryos. Molecular biology of the cell. 2006;17(11):4911–4924.

62. Canman JC, Hoffman DB, Salmon ED. The role of pre- and post-anaphase microtubules in the cytokinesis phase of the cell cycle. Curr Biol. May 18 2000;10(10):611–4. doi:10.1016/s0960-9822(00)00490-5

63. Schindelin J, Arganda-Carreras I, Frise E, et al. Fiji: an open-source platform for biological-image analysis. Nature Methods. 2012/07/01 2012;9(7):676–682. doi:10.1038/nmeth.2019

64. Dobin A, Davis CA, Schlesinger F, et al. STAR: ultrafast universal RNA-seq aligner. Bioinformatics. Jan 1 2013;29(1):15–21. doi:10.1093/bioinformatics/bts635

65. Liao Y, Smyth GK, Shi W. featureCounts: an efficient general purpose program for assigning sequence reads to genomic features. Bioinformatics. 2013;30(7):923–930. doi:10.1093/bioinformatics/btt656

66. Love MI, Huber W, Anders S. Moderated estimation of fold change and dispersion for RNA-seq data with DESeq2. Genome Biol. 2014;15(12):550. doi:10.1186/s13059-014-0550-8

