## Supplementary figures and images for "The Unkempt RNA binding protein reveals a local translation program in centriole overduplication"

### Supplemental Figure 1

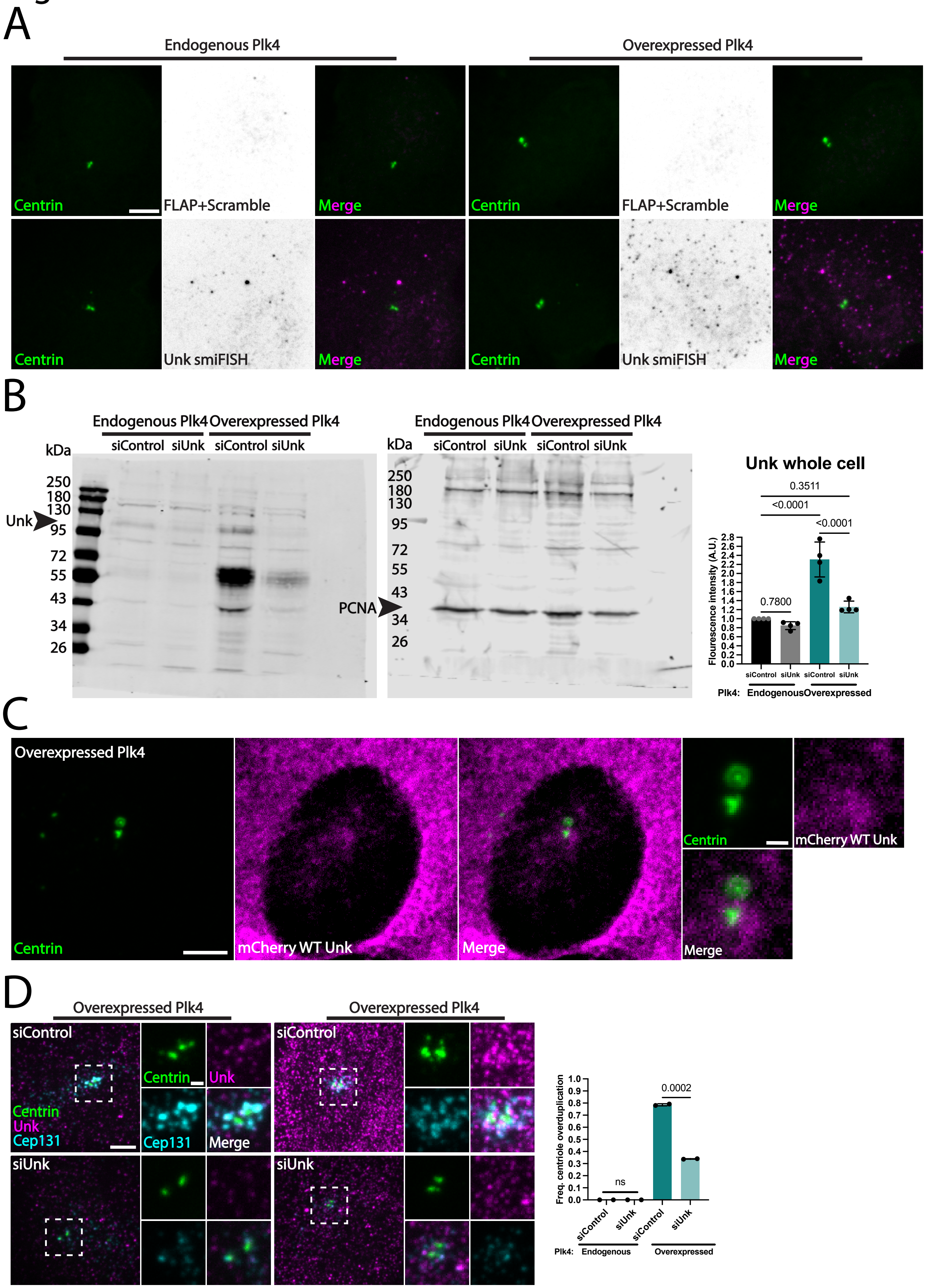

### Supplemental Figure 2

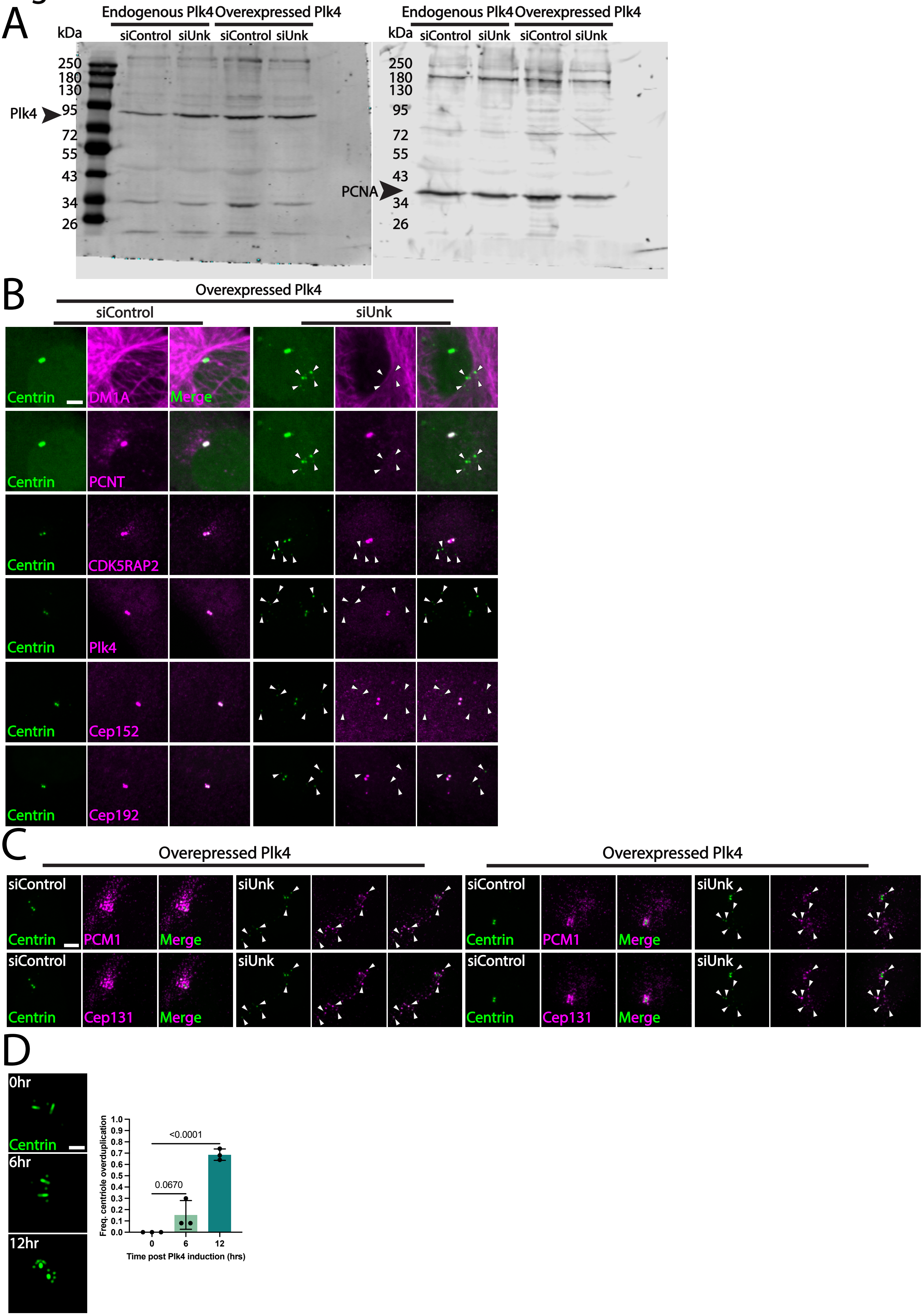

### Supplemental Figure 3

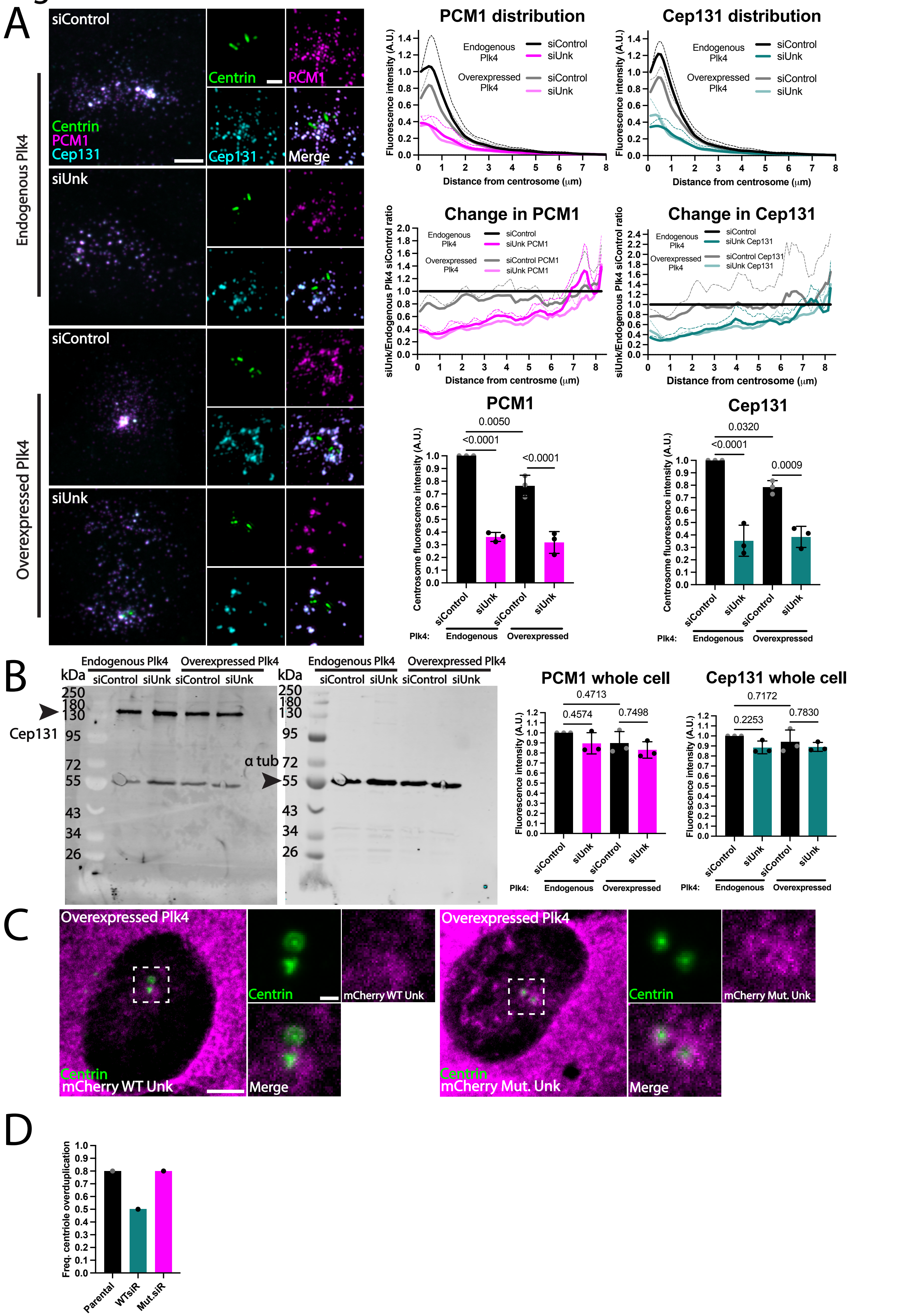

### Supplemental Figure 4

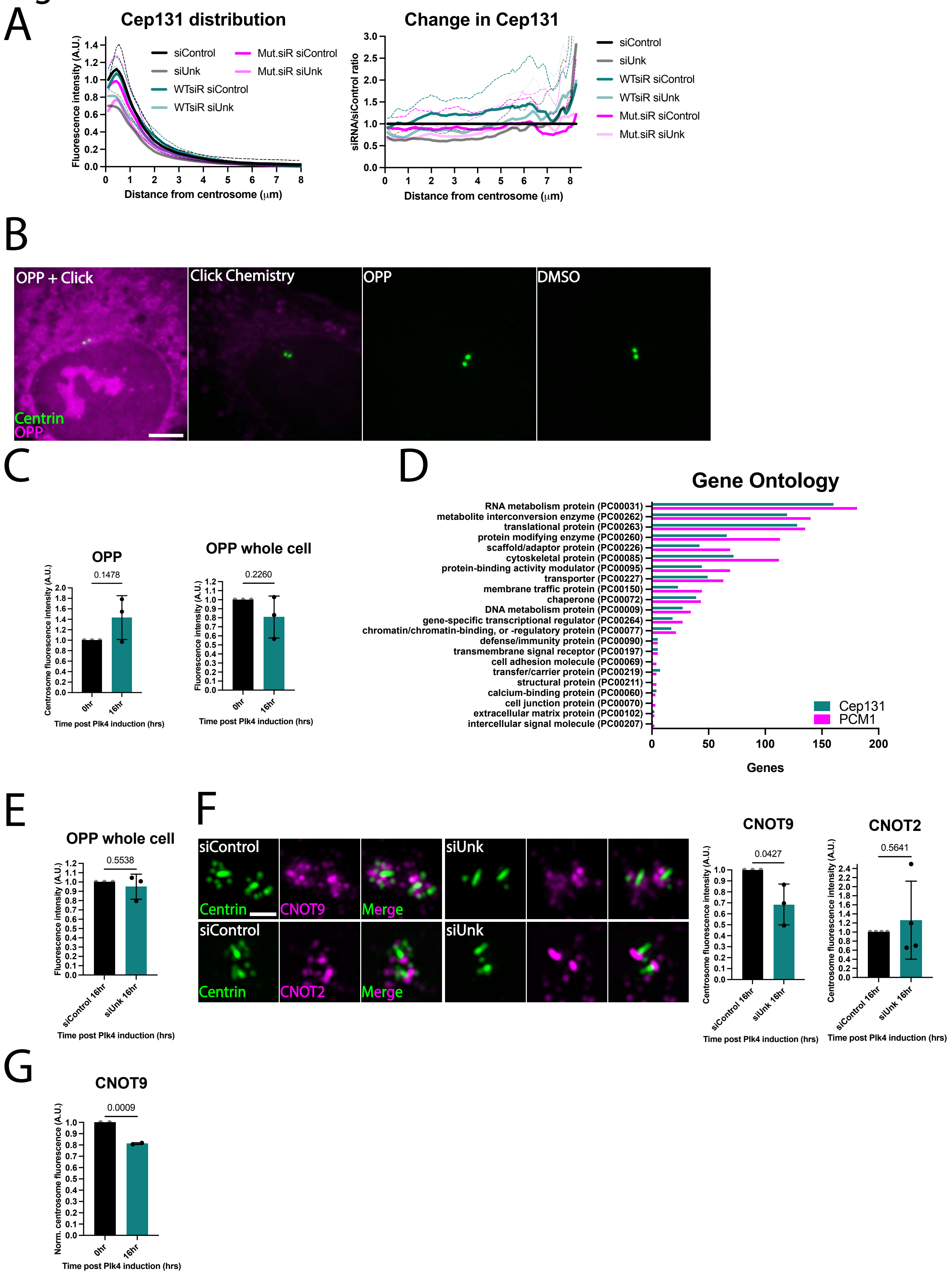
